# Development of long-term oyster tissue cultures reveals cellular plasticity, regenerative potential, and sustained physiological activity

**DOI:** 10.1101/2025.11.07.687009

**Authors:** Kallen R. Sullivan, Pieter C. Steketee, Scott Maxwell, Liam J. Morrison, Tim Regan, Tim P. Bean

**Affiliations:** The Roslin Institute, University of Edinburgh, EH25 9 RG, United Kingdom; The Royal (Dick) School of Veterinary Studies, University of Edinburgh, EH25 9 RG, United Kingdom

## Abstract

Molluscs are ecologically and economically significant, yet progress in understanding their cellular biology has been constrained by the lack of reliable molluscan cell culture systems. Insights into key molluscan biology such as immunity, biomineralization, and host-pathogen interactions, has previously been limited by short term survival of primary culture systems. Here, we establish reproducible long-term *in vitro* and *ex vivo* cultures of cells and tissues from the oyster, *Ostrea edulis*, including whole heart, visceral mass, mantle, and gill explants. Explanted tissues remained viable for over two months, with whole hearts maintaining rhythmic, stimulus-responsive contractions for up to nine months. Cultures maintained structural and functional features while uncovering cellular plasticity, including haemocyte granule turnover, auricular epithelial remodelling with novel ciliation, and the emergence of *de novo* multicellular microtissues. Previously under-characterized tissue-specific cell types were maintained in culture, including adipocytes from the visceral mass and myocytes from the mantle, thereby expanding the cellular repertoire accessible *in vitro*. Mineralized structures were observed across tissue cultures, highlighting the potential of this system for studying biomineralization. By sustaining diverse cell types *in vitro*, these cultures provide a novel tractable platform for studying molluscan diseases, while also laying the foundation for deeper investigation into comparative invertebrate biology and the development of immortalised molluscan cell lines.

## Introduction

Cell culture systems provide a critical bridge between organism-level biology and the underlaying cellular mechanisms. Experimental *in vitro* systems enable detailed exploration of cell physiology, behaviour and disease modelling, thereby progressing our biological understanding. For molluscs, whose ecological functions and aquaculture value are globally significant, the absence of stable, reproducible culture systems has hindered progress in studies ranging from immunity and stress physiology to biomineralization and disease management.

Molluscs are among the most diverse and ecologically significant animal phyla, comprising more than 86,000 described species and likely exceeding 200,000 across marine, freshwater, and terrestrial ecosystems ^1^. They occupy critical ecological roles while also underpinning global aquaculture, fisheries, and biotechnological industries ^2^. Oysters, in particular, highlight this importance: global aquaculture production is approximately 5.5 million tonnes, and they provide ecosystem services such as water filtration, sediment stabilization, and reef formation that enhance coastal biodiversity ^3,4^. Beyond industry and ecology, molluscs are established models in neurobiology, biomineralization, and ecotoxicology ^5^. Yet, despite this breadth of significance, fundamental insights into their cellular biology remain limited, constrained by persistent challenges in establishing reliable primary cell cultures ^6^.

Attempts to develop molluscan cell cultures date back several decades ^6,7^. Haemocytes, the circulating immune cells of molluscs, are the most frequently studied cell type because they are readily accessible and play a central role in immune response and wound healing ^8^. Thus, often used in functional immune assays, which typically run over hours to a few days; in this context, their limited viability does not preclude utility. However, their limited lifespan and terminal differentiation restrict investigations into long-term processes, including biomineralization, developmental biology, chronic stressors, and neoplastic transformation ^9–11^. Moreover, reliance on repeated haemocyte isolations introduces a higher degree of animal-to-animal variability that limits reproducibility and scalability ^12,13^.

To date, only two molluscan systems have been propagated beyond short-term primary culture. The *Biomphalaria glabrata* embryonic (Bge) line, derived from snail embryos, a continuous freshwater mollusc line, maintained since the 1970s and widely used to study host-parasite interactions with *Schistosoma mansoni* and invertebrate immunity ^14,15^. More recently, the *Chlamys farreri* Trochophore (CfT) line, derived from Zhikong scallop trochophore larvae, was reported as the first long-term scallop culture to be established ^16^. However, its designation as a continuous line will remain unresolved until key parameters, including long-term stability across subcultures are demonstrated, and the cellular composition is completely defined. Together, Bge and CfT demonstrate that molluscan cells can be maintained in extended culture, but they also highlight unresolved challenges: extremely limited taxonomic representation, a reliance on embryonic or larval sources that may not capture adult tissue physiology, and incomplete validation in terms of cell identity, passage stability, and reproducibility. Ultimately, immortalisation is not an end in itself; the value of any molluscan cell line will depend on whether it can support targeted application, provides a reproducible model for biological study, and it is genetic tractable. These challenges highlight both the technical barriers that remain and the transformative potential that could be realised if they are overcome.

Precedent from insects provide a clear success story for invertebrates: cell lines such as *Drosophila Schneider 2* and *Spodoptera frugiperda clone 9* have become indispensable tools in virology, immunology, and recombinant protein production ^17,18^. Recently, the first continuous marine invertebrate cell line was established from the sponge *Axinella donnani* ^19^. Its development required customized media adapted to the osmotic and nutritional demands of sponge cells, underscoring the need to align culture conditions with the unique physiology of marine invertebrates. This achievement demonstrates that stable marine invertebrate cell lines are attainable, marking a clear path forward for similar advances in marine molluscs.

To address the limited longevity of haemocyte cultures and limits of immortalised lines for marine molluscs, explanted tissues have been employed as an alternative means of capturing and maintaining molluscan cells *in vitro*. In this context, “explants” refer to small fragments of tissue maintained under culture conditions, from which cells can migrate, proliferate locally, or remain organized within the tissue. In contrast to dispersed haemocyte suspensions, explants preserve aspects of the native architecture, including cell-cell and cell-matrix interactions, which can sustain viability and functional integrity over longer intervals ^6,7^. Direct evidence for extracellular matrix secretion by explanted molluscan tissues is limited. However, studies report that the addition of tissue extracts can improve culture yield and adherence, indicating that tissue-derived factors may influence the culture environment ^20–22^. It is therefore likely that explants contribute to culture medium conditioning, which could partly alleviate the mismatch between mammalian-formulated media and the physiological requirements of marine invertebrate cells. Explants also provide access to a broader array of tissue-resident cell types, creating opportunities to study cellular diversity. Cells derived from mantle, gill, digestive gland, and gonadal tissues have most commonly been maintained *in vitro* with survival periods ranging from several days to several months ^23–26^. These systems extend culture longevity and broaden the range of cell types available; however, achieving longer-term primary culture is critical for fully characterizing molluscan cells and represents a necessary step toward developing immortalized, standardized lines.

Among bivalves, the European flat oyster, *Ostrea edulis*, is a high-priority species. It supports valuable aquaculture yet remains highly vulnerable to pathogens such as *Bonamia* spp., which continue to cause recurrent mortality events ^27,28^. Beyond its commercial significance, *O. edulis* has become a flagship for large-scale restoration initiatives across Europe, where the recovery of lost oyster reefs is central to rebuilding coastal biodiversity and ecosystem services ^29,30^.

Recognizing this strategic importance, we developed optimized culture conditions that support the long-term maintenance of multiple *O. edulis* tissues. The primary aim of this study was to establish and characterize long-term *in vitro* and *ex vivo* cultures of oyster cells and tissues. To achieve this, our specific objectives were to (i) evaluate the longevity of mantle, gill, adductor muscle, visceral mass (VM), and heart explants under uniform conditions, (ii) assess tissue-level organization and physiological activities such as contractility and ciliation, and (iii) characterize the diversity and behaviour of resident and shed cell populations.

Our cultured mantle, gill, VM, and whole-heart explants revealed diverse cell populations, including haemocytes, fibroblasts, myocytes, ciliated epithelial cells, and adipocytes, along with novel phenomena including epithelial ciliation, haemocyte granule plasticity, and spontaneous microtissue formation. Together, these cultures provide new opportunities to explore molluscan cellular diversity and to develop more sophisticated, long-term model systems.

## Methods

### Oyster sourcing and storage

Adult European flat oysters (*Ostrea edulis*; small-medium class, >6.5cm) were obtained from a commercial supplier in the United Kingdom and shipped overnight in insulated containers. For all procedures except haemolymph cytology, oysters were either dissected immediately upon arrival or stored at 4 °C and processed within two days.

For haemolymph cytology preparations, oysters were instead transferred on arrival to aerated aquaria containing artificial seawater (ASW; 35 ppt). Animals were maintained at 16 °C without feeding for one week to allow depuration, with water changed every other day before haemolymph extraction.

### Artificial seawater

ASW was prepared by dissolving Tropic Marin Classic Sea Salt (Wartenberg, Germany) in Milli-Q purified water at two concentrations: 35 g/L (35 ppt) for oyster maintenance and tissue wash/holding, or 60 g/L (60 ppt) for media preparation. All ASW solutions were sterilized by autoclaving.

### Tissue dissections and explant yield

Dissections were performed by inserting a shucking knife at the hinge, then severing the adductor muscle, and removing the right valve (Fig 1A,B). The VM was separated along the transverse plane just below the pericardium using a scalpel to improve visibility of the heart (Fig 1B). The whole heart was excised by removing the pericardium and tracing the margins of the auricles and ventricle with spring scissors (Fig 1B, 1C). The VM was then bisected along the coronal plane to create two equal halves, shaved to a thickness of 4-6 mm, and sampled with a 6 mm² biopsy punch (Fig 1B, 1C). Individual layers of the mantle and gill were isolated for sampling. Mantle tissue was sampled by centring a 4 mm² biopsy punch on the edge of the lobes, producing half-moon-shaped explants (Fig 1B, 1C). Individual gill layers were gently stretched to ensure even tension before sampling with a biopsy punch; this yielded uniformly round explants at the time of sampling, though the natural grooves of unstretched gill tissue later caused explants to appear oval in culture (Fig 1B, 1C). The adductor muscle was bisected along the coronal plane, trimmed to a thickness of 4-6 mm, and sampled with a 6 mm² biopsy punch (Fig 1B, 1C).

**Fig 1.**
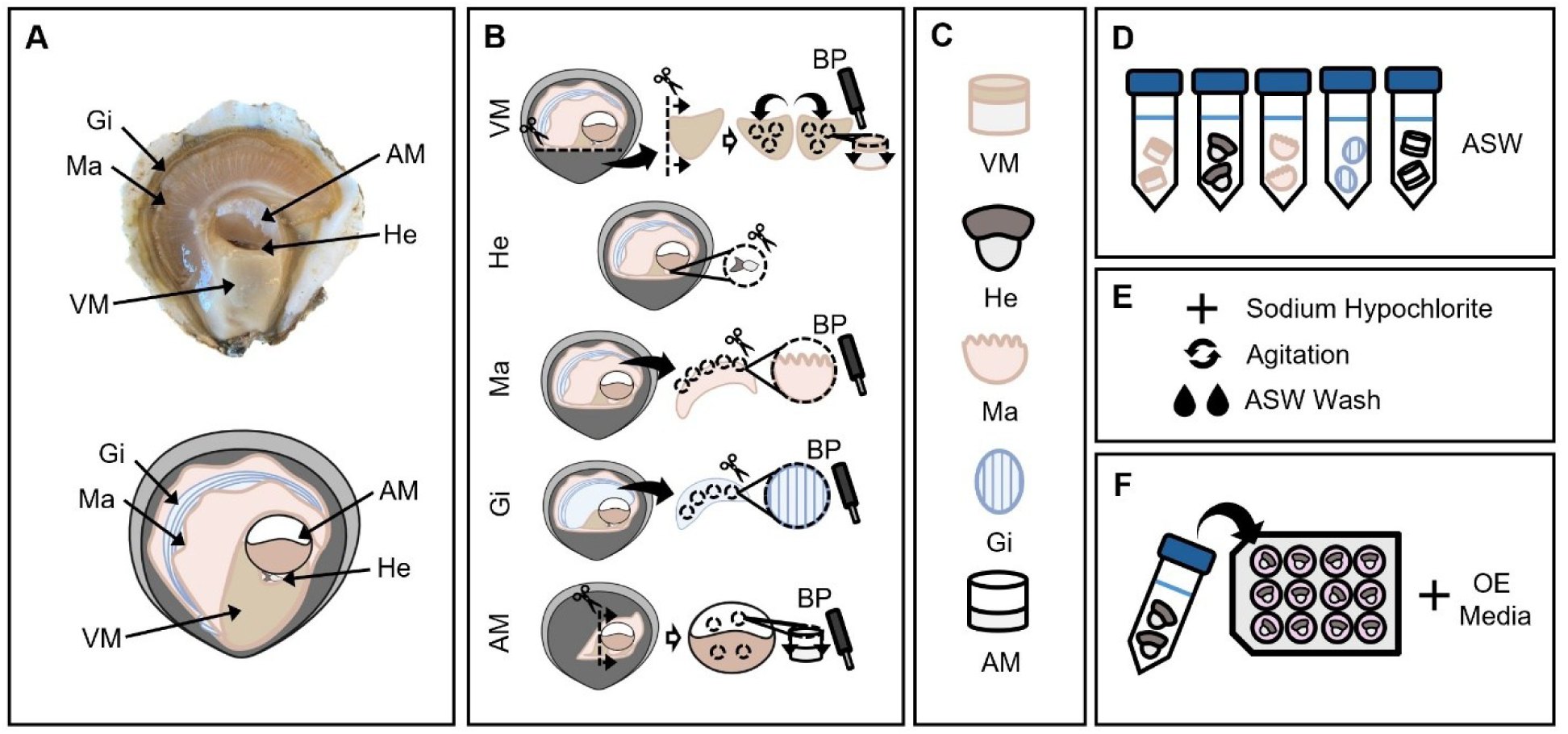
Workflow for *Ostrea edulis* tissue explant preparation and culture. **(A)** Schematic of oyster anatomy**. (B)** Tissue collection: visceral mass, heart, mantle, gill, and adductor muscle were dissected from whole animals and sectioned into tissue explants using spring scissors or a biopsy punch. **(C)** Representative icons depicting the general morphology of each tissue explant. **(D)** Explants were pooled by tissue type and placed into tubes containing artificial seawater. **(E)** Explants were surface sterilized with sodium hypochlorite, agitated, and washed twice with artificial seawater. **(F)** Sterilized explants were transferred into culture plates with *Ostrea edulis* (OE) media for downstream experiments. Abbreviations: AM, adductor muscle; ASW, artificial seawater; BP, biopsy punch; GI, gill; HE, heart; MA, mantle; OE, Ostrea edulis; VM, visceral mass.

Dissections from a single adult oyster yielded one whole heart explant, approximately 6-8 digestive gland explants, 10-12 mantle explants, 12-16 gill explants, and 4-6 adductor muscle explants. All dissected tissues were pooled by tissue type and placed in 50 mL Falcon tubes containing 35 ppt ASW (Fig 1D).

### Culture medium

*O. edulis* (OE) medium was prepared by mixing equal parts of L-15 (Leibovitz) medium (Merck) and 60 ppt ASW, resulting in an osmolality of approximately 950 mOsm/kg, as measured using an Advanced Model 3320 Micro-Osmometer (Advanced Instruments). This formulation was adapted from *Potts et al.* (2020) but adjusted to match *O. edulis* haemolymph osmolality (940 ± 9 mOsm/kg, n=4) and to include a lower concentration of Gibco Antibiotic-Antimycotic (0.1%). The final mixture was passaged through a 0.45 µm filter before use.

### Decontamination, culture conditions and monitoring

Following dissection, tissue explants underwent a decontamination process. Samples were treated with a 0.05% (v/v) active sodium hypochlorite solution for 10 minutes at room temperature, with occasional agitation to improve surface sterilization. This was followed by two successive rinses in 35 ppt ASW, each lasting 10 minutes in a sterile tissue culture hood (Fig 1E). After disinfection, tissue pieces were aseptically transferred to 6-, 12-, or 24-well culture plates containing the *O. edulis* culture medium with 3 mL, 2 mL or 1 mL of OE media, respectively (Fig 1F). Plates were incubated in the dark at 20 °C. After 24 hours, cultures were checked for any obvious signs of microbial contamination before being transferred to fresh wells and media.

Negative biological controls were prepared using either low-salinity (650 mOsm/kg) or high-salinity (1100 mOsm kg) media. Tissues in low-salinity media served as models to observe hypoosmotic stress, whereas those in high-salinity media were used to assess apoptotic features such as cell fragmentation and loss of adherence. Wells containing only media were maintained throughout the culture period as contamination controls. Tissues were maintained in culture and monitored regularly for signs of dissociation, microbial contamination (e.g. bacteria, protozoa), or the accumulation of shed apoptotic cells. Explants showing evidence of microbial contamination were immediately discarded to prevent cross-contamination.

### Live cell characterisation

Cells shed into the media were routinely examined across tissue culture types. Qualitative observations were documented throughout the culture period, focusing on cell presence, relative abundance, morphology, and behaviour.

For cytological staining, cells shed into the media (excluding non-adherent adipocytes from visceral mass explants) and fresh haemolymph were processed with the Hemacolor rapid staining system (Merck). For cells shed into culture media, 20 µL of culture medium was pipetted onto Superfrost Plus slides (Thermo Fisher Scientific), air-dried, and fixed in ice-cold 100% methanol for 7 min. For fresh haemolymph, samples were withdrawn from the adductor muscle using a sterile 18-gauge needle attached to a 1 mL syringe, placed on Superfrost Plus slides (Thermofischer), and mixed 1:1 with 35 ppt ASW to promote cell spreading. The mixture was left for 1 hour at room temperature to allow haemocytes to settle, then excess fluid and debris were gently removed with ASW. Slides were air-dried and fixed in ice-cold methanol for 7 min. In both cases, fixed slides were stained according to the manufacturer’s protocol, mounted with DPX medium (Sigma-Aldrich), and sealed with coverslips.

### Tissue and microtissue characterisation

Fresh, 3-month cultured, and 6-month cultured oyster tissues processed directly from the culture medium by the Histopathology Laboratory, Diagnostic Biopsy Service, The Royal (Dick) School of Veterinary Studies, University of Edinburgh, following standard operating procedures. Samples were fixed in neutral buffered formalin, embedded in Histoplast IM paraffin wax (Epredia), and sectioned at 4 µm using a HM340E microtome (Epredia). Sections were mounted on Slidemate Plus adhesion microscope slides (Epredia) and coverslipped with Pertex mounting medium (CellPath).

Microtissues recovered from culture media and isolated adipocytes were first embedded in Histogel (Epredia) prior to sectioning and were processed in parallel with paraffin-embedded tissues.

To assess tissue architecture and cellular features, standard histological stains were applied, including haematoxylin and eosin (H&E), Masson’s Trichrome, Masson’s Trichrome with Alcian Blue counterstain, Masson’s Fontana, Perls’ Prussian Blue, and Rhodanine. Staining was performed primarily using a Gemini Varistain autostainer (Thermo Fisher Scientific). When staining intensity was suboptimal, slides were restained manually to ensure consistent contrast and interpretation.

Prepared slides were scanned at ×40 magnification using a Hamamatsu NanoZoomer XR slide scanner. Digital images were archived in NDPI format and analysed with NDP.view2 software (Hamamatsu Photonics).

### Assessment of heart rate during *ex vivo* culture

The heart rates of five heart explants were monitored to assess baseline (resting) and post-stimulation (media change) activity. Explants were maintained in culture for a total of 35 days, spanning 5 full weeks of culture, and heart rate measurements were taken once weekly (day 0, 7, 14, 21, 28, 35). During each observation session, videos of beating heart explants were captured. Recordings were subsequently replayed for manual analysis, during which the number of visible contractions was counted over a 2-minute interval and divided by two to calculate beats per minute (BPM). Heart explants were recorded both immediately before and immediately after media replacement to compare resting and stimulated activity.

A two-way repeated measures ANOVA was performed with Time (six observation points) and Media Replacement Status (pre-vs. post-media change) as within-subjects factors. Where significant main effects were observed, Tukey’s post hoc tests were conducted to identify pairwise differences. Statistical significance was defined as *p* < 0.05.

### Image analysis and cell measurements

Microscopy was performed using multiple systems according to application. Histological slides were imaged on a Nikon ECLIPSE Ni upright microscope with 10×, 20×, and 40× air objectives and a 100× oil-immersion objective, using a ZEISS Axiocam 105 colour camera and ZEN Blue software. Microtissue time-lapse imaging was carried out on a Zeiss Axio Observer inverted microscope equipped with an environmental chamber (20 °C, no CO₂) and a 20× objective, with images acquired on a ZEISS Axiocam 807 mono camera through ZEN Blue Pro with automated acquisition every 15 minutes. Manual time-lapse and live-cell phase-contrast imaging were performed on a Zeiss Axiovert 40 inverted microscope with a ZEISS Axiocam MRm camera and ZEN 2 software with manual acquisition every 30 seconds. Additional live-cell imaging, including microtissue and heart rate recordings, was obtained using a Leica DMi1 inverted microscope with Leica imaging software.

Following image acquisition, standard adjustments (rotation, flipping, and cropping) were applied in ImageJ (NIH, version 1.54). Digitized histological slides were analysed in NDP.view2 software, while cytology and live-cell images were measured in ImageJ. For each cell type or morphological feature, 100 measurements were collected to generate representative size distributions, except for hyaline globules, where 200 measurements were taken. Fields of view were randomly selected and all cells counted to minimize sampling bias. Results are reported as mean ± standard deviation (SD) with observed ranges.

## Results

Improved culture media supported the long-term maintenance of multiple oyster tissues and provides a platform for *in vitro* studies of tissue and cellular behaviour. Maintenance was defined as preservation of intact structure and activity (e.g. heart contractility) without dissociation. Using this approach, cultures were sustained for extended periods: mantle up to 4 months, gill up to 2 months, VM for 8.5 months, and contracting hearts up to 9 months. During culture, two distinct cell populations were monitored: cells shed into the medium and cells retained within the explants. Shed cells included haemocytes, myocytes, fibroblasts, ciliated epithelial cells, and adipocytes, most of which could be traced to their counterparts in the explants.

### *Ex vivo* culture of intact hearts

Whole-heart cultures were established by transecting the veins connecting the ventricle and auricle while preserving the surrounding tissue (Fig 2A). *In situ*, hearts displayed an elongated, stretched conformation, but in culture they adopted a more compact morphology (Fig 2A-B). Cultured hearts retained rhythmic contractility for up to nine months and remained responsive to external stimuli, with beat frequency increasing immediately after media changes.

**Fig 2.**
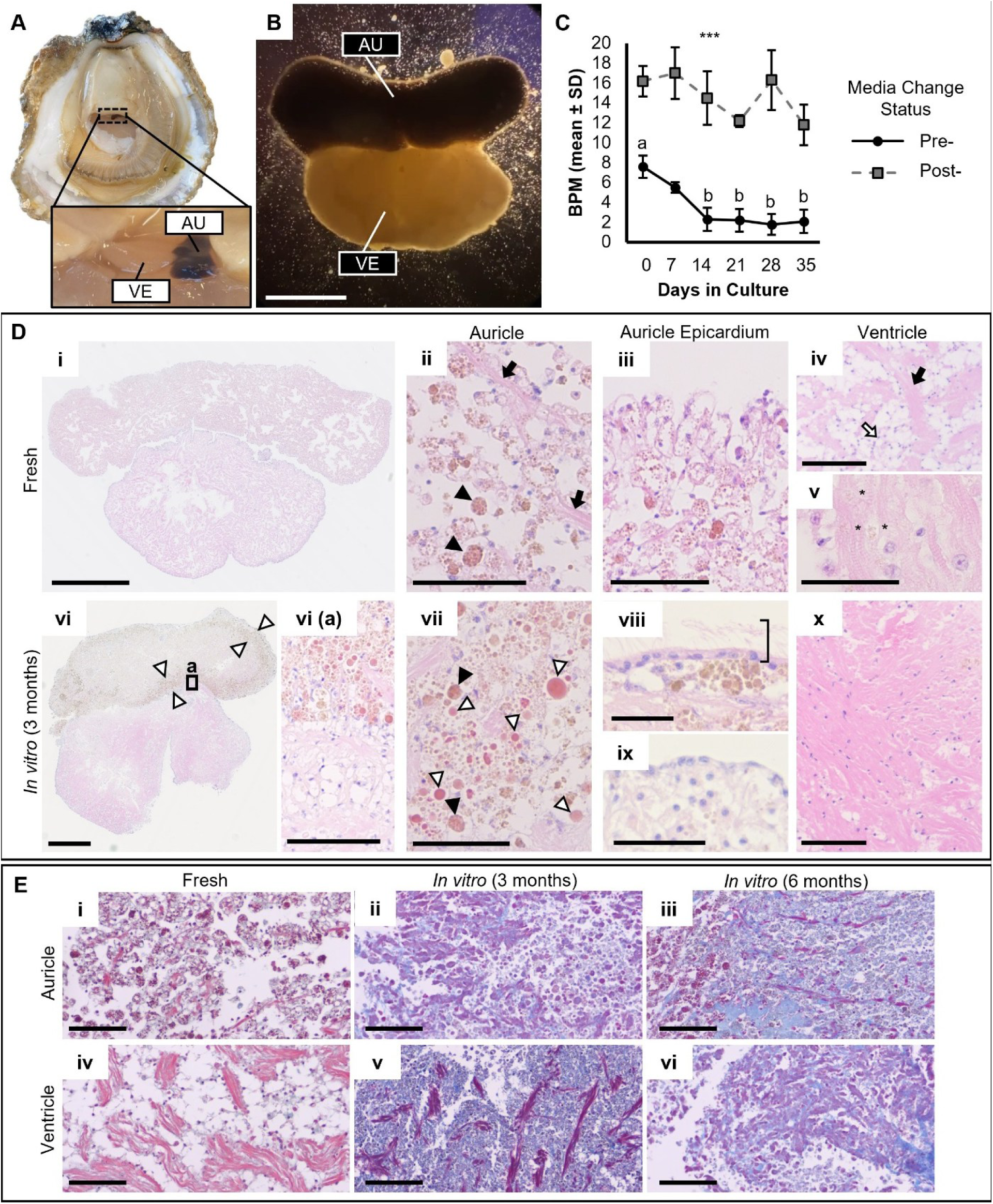
Morphology, function and histological changes of *Ostrea edulis* hearts during long-term culture. **(A)** Image of a whole oyster with the right valve removed. Inset shows an intact heart with VM and pericardium removed, exposing the auricle (AU) and ventricle (VE). **(B)** Image of an isolated heart explant in culture, showing distinct auricle (AU) and ventricle (VE) regions. **(C)** Mean ± SD beat rates measured before (pre-) and after (post-) media change at 6 timepoints in culture (n = 5 explants). Post-media change rates were consistently higher than pre-media change rates (***, p < 0.001). Letters denote significant post-hoc differences in pre-media change rates: Day 0 (a) was significantly higher than days 14 - 35 (b), p < 0.02. **(D)** Histological sections of fresh (i - v) and 3-month cultured (vi - x) hearts stained with H&E. Representative features are indicated: granule clusters (closed arrowheads; ii, vii), cardiomyocyte bundles (closed arrows; ii, iv), fibroblast-like connective tissue (open arrows, iv), lipofuscin deposits (asterisks, v), region of localised concentration of hyaline globules (paired open arrowheads; vi), individual hyaline globules (open arrowheads, vii), and ciliated auricular epicardium (bracket, viii). **(E)** Histological sections of fresh tissue (i, iv), 3-month (ii, v) and 6-month (iii,vi) cultured hearts stained with Masson’s Trichrome showing progression of collagen deposition (fibrosis) overtime. Scale bars: (B, D(i), D(vi)) = 2 mm; (D(ii-iii), D(vii-ix)) = 50 µm; (D(iv), D(x), D(vi)(a), E(i-vi)) = 100 µm.

To assess the acclimatization time required for hearts to adjust to culture conditions, contraction rates were monitored weekly for five full weeks of culture, both before and after media replacement (Fig 2C; S1 Table). A two-way repeated measures ANOVA showed a significant main effect of culture time, F(5,20) = 3.09, p = 0.032, and of condition (pre- vs. post-media change), F(1,4) = 144.05, p < 0.001, with no significant interaction, F(5,20) = 1.23, p = 0.332. Thus, beat rates varied across weeks and were consistently higher immediately after media replacement compared with pre-change values (Fig 2C). Pre-media change beat rates declined significantly over time, F(5,20) = 7.45, p < 0.001. Tukey post hoc tests indicated that day 0 rates were higher than day 14 - 35 (all p < .02), whereas rates between day 7 to 35 did not differ significantly (Fig 2C). In contrast, post-media change beat rates remained stable across the culture period, F(5,20) z= 1.31, p = 0.300, suggesting that the immediate contractile response to fresh media was maintained over time. These findings demonstrate that oyster hearts retain contractile activity and responsiveness for extended periods *in vitro* and indicate that 14 days in culture is sufficient for acclimatization. After 14 days *in vitro*, the average heart rate of explants before media replacement was 1.82 ± 2.09 bpm, whereas after media replacement it increased to 13.90 ± 5.10 bpm.

Functional stability was accompanied by preservation of tissue architecture. In freshly dissected hearts, auricles and ventricle displayed distinct cellular compositions (Fig 2D(i)). Auricles were dominated by granule-containing connective tissue cells (Fig 2D(ii)) and lined by a wavy squamous epicardium, likely reflecting loss of tension after dissection (Fig 2D(iii)). The ventricle consisted mainly of cardiomyocytes with scattered connective tissue (Fig 2D(iv)).

After three months in culture, hearts were reduced in size and cells appeared more densely packed (Fig 2D(vi)), yet clear boundaries between auricles and ventricle were maintained (Fig 2D(vi)(a)). No obvious signs of necrosis or central cell loss was observed by histology, indicating sustained tissue viability. Histological analysis revealed thickening of the epicardium, with most of the auricular surface acquiring motile cilia (Fig 2D(viii); S1 Video), although some regions remained cilia-free (Fig 2D(ix)). Ciliated cells appeared consistently between three and six months, suggesting a temporal progression of epithelial transformation under culture conditions. The epicardial layer also shifted from a squamous to a columnar-like morphology, with nuclei aligned along the basement membrane of the ciliated cells (Fig 2D(viii)). Ventricle tissue became increasingly compact, with a reduction in connective tissue cells between the cardiomyocytes and haemolymph sinuses (Fig 2D(x)).

Auricle tissue was almost entirely dominated by granule-containing connective-tissue cells; individual cells with distinctive membrane-bound cytoplasmic inclusion which occasionally aggregate into larger dense packages (Fig 2D(vii)). These were uniformly distributed throughout the auricle but absent from ventricle tissue in both fresh and cultured samples (Fig 2D(iv), 2D(x)). Histochemical staining identified the granules as PAS-positive, indicating carbohydrate-containing material. On H&E they appeared dark pink to brown, and with Masson’s Trichrome they stained dark pink, suggesting a proteinaceous component. No staining was detected with Perls’ Prussian Blue, Masson’s Fontana, or Toluidine Blue, excluding iron, melanin, and mast cell proteoglycans. The granules were also distinct from lipofuscin, which typically occurs as small, irregularly distributed pigment deposits (Fig 2D(v)). In contrast, granules in auricular cells were larger, uniformly distributed, and present in both fresh and cultured tissues, excluding age-related pigment accumulation as their identity. Granules from fresh tissue measured an average of 1.81= ± 0.63 µm in diameter (0.82 - 3.55 µm; S2 Table), while larger aggregates consisting of multiple membrane-bound granules averaged 11.32 ± 1.90 µm (7.87 - 17.63 µm; S2 Table; Fig 2D(vii)). In addition to the intracellular granules, extracellular deposits were observed in long-term cultures. At the three-month sampling point, a distinct ring of hyaline globules was present along the auricular perimeter at the ventricular margins or adjacent to newly formed ciliated cells (Fig 2D(vi)). These globules measured an average of 5.14 ± 1.60 µm in diameter (2.55 - 11.40 µm; S2 Table). At six months, the hearts displayed the same prominent ring of hyaline globules and ciliation observed in earlier cultures, but the surrounding tissue showed progressive collagen accumulation, with the six-month cultures appearing notably more fibrotic than fresh or three-month cultured hearts (Fig 2E(i-vi)).

When auricle tissue was cultured independently, the majority of shed cells consisted of the granule-containing connective-tissue cells, consistent with their abundance in this region (Fig 2D(ii),2D(vii); 3A). In culture, these cells were non-adherent and measured 14.84 ± 1.77 µm in diameter (range: 11.26-19.14 µm; S2 Table). A subset displayed a concentric morphology, with a spherical outer boundary enclosing a single densely pigmented internal body (Fig 3B). In some cases, three or more internal bodies were present within one cell (Fig 3C, 3D). The relationship of these additional structures to the granule aggregates observed in parent tissue remains unresolved.

**Fig 3.**
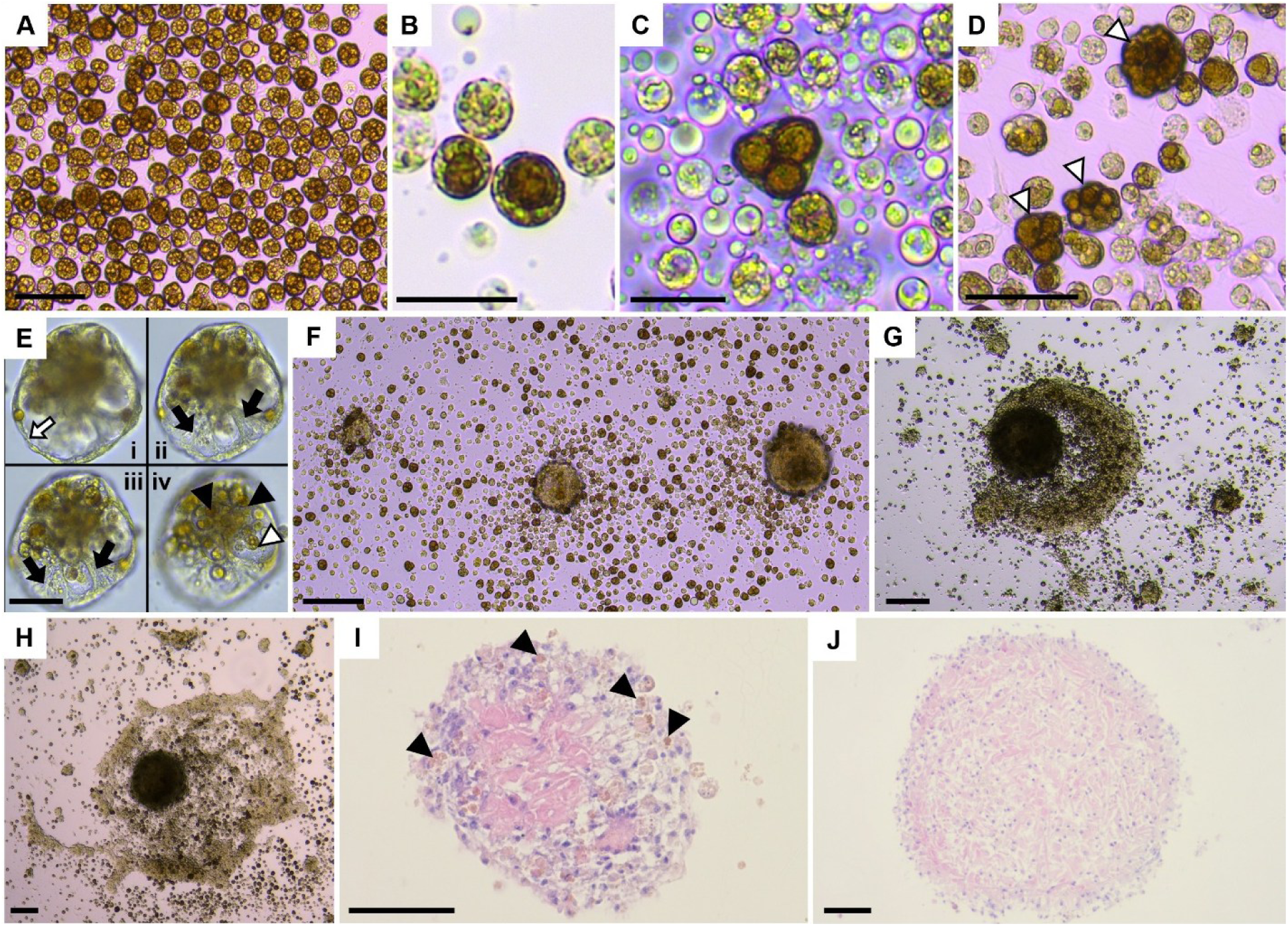
Auricle-derived cells and emergence of contractile microtissues in cultured hearts of *Ostrea edulis*. **(A)** Granule cells derived from auricle-only culture (ventricle removed). **(B)** Auricle-derived cells with concentric morphology. **(C-D)** Auricle-derived cells containing multiple internal bodies (open arrowheads). **(E)** Contracting microtissue. Sequential focal planes (i - iv) progress from the top surface (i) to the bottom adjacent to the culture plate (iv). Auricle-associated granule cells (closed arrowhead), auricle-associated granule cells containing multiple internal bodies (open arrowhead), contractile regions (closed arrow), and tissue margins (open arrow) are indicated. **(F)** Representative field of view showing three early microtissues, ranging from a small adherent structure (left) to larger microtissues with upward growth and defined boundaries (right). **(G-H)** Large contractile microtissue with concentric ring-like outgrowths. **(I)** H&E-stained microtissue fixed in Histogel, containing auricle-derived granule cell type. **(J)** H&E-stained microtissue fixed in Histogel, composed predominantly of cardiomyocytes. Scale bars: (A, D, E(i-iv)) = 50 µm; (B, C) = 20 µm; (F-J) = 100 µm.

In addition to individual cells, organized multicellular structures were frequently observed as distinct from explants in long-term heart cultures (Fig 3E-H). These microtissues typically began as small semi-adherent clusters that enlarged and detached into suspension, where they exhibited contractions and dispersed evenly across the culture dish (Fig 3E-F; S2 Video). Larger microtissues occasionally resettled and developed concentric ring-like outgrowths (Fig 3G-H). Unlike loosely formed floating aggregates, these masses displayed clear structural boundaries, indicating a higher degree of cellular organization. Live-cell time-lapse imaging demonstrated division and outward growth of an auricle-derived cell will multiple internal bodies (S3 Video). Pigmented granule cells in culture shared features with the granule-containing auricle cells described in intact tissue, indicating continuity between tissue-resident and shed populations. Some microtissues expanded outward from a central core, resembling the subset of granule cells containing three or more internal bodies rather than forming through simple cell aggregation (Fig 3C; S3 Video). Histological analysis confirmed that the *de novo* microtissues were comprised of cell types present in the parent tissue, including auricle-associated granule cells and cardiomyocytes (Fig 3I), with some consisting predominantly of cardiomyocytes (Fig 3J). Overtime, a subset of microtissues also developed a ciliated phenotype, consistent with the ciliation observed in the auricle region of long-term whole-heart cultures (S2 Video).

### Dynamic behaviour of haemocytes *in vitro*

Haemocytes freshly isolated from haemolymph exhibited two primary morphologies under phase contrast live cell imaging. The first population, consistent with hyalinocytes, had uniformly dark cytoplasm with elongated, fibroblast-like adherence and fine filopodia extending from the cell body; these cells often overlapped extensively when interacting with neighbours (Fig 4A-B). The second population, consistent with granulocytes, appeared more rounded when fully adhered, with clearly defined central nuclei and heterogeneous cytoplasmic granulation (Fig 4C-F). Under phase-contrast microscopy, granules appeared as large, bright, pearl-like inclusions, surrounded by a halo of fine dark granulation concentrated in the central region of the cell body (Fig 4C, 4F; S4 Video). These cells typically aggregate margin-to-margin, maintaining more distinct boundaries while clustering in groups (Fig 4C). Although both morphologies were present in fresh haemolymph, only the rounded granulocyte type was consistently maintained in explant cultures across all tissue types (Fig 4D-F). Staining of fresh haemolymph further revealed additional heterogeneity, with multiple cell populations or states distinguishable by their degree of granulation (Fig 4G-K). Cells shed into explant cultures closely resembled the following haemolymph phenotypes: some contained small refractive eosinophilic inclusions (cf. Fig 4H, open arrowhead), others had larger basophilic inclusions (cf. Fig 4J-K), and some lacked inclusions altogether (cf. Fig 4G, double plus). In heart cultures, the “true” hyalinocyte population were observed only during the initial days of culture but did not persist (cf. Fig 4A). In addition to differences in granulation and morphology, some haemocytes displayed variation in nuclei ranging from bilobed and multilobed forms in both fresh haemolymph and cultured haemocytes (Fig 4L-N). These lobed nuclei were less frequent than typical rounded forms and did not appear to be to be restricted to a particular haemocyte type.

**Fig 4.**
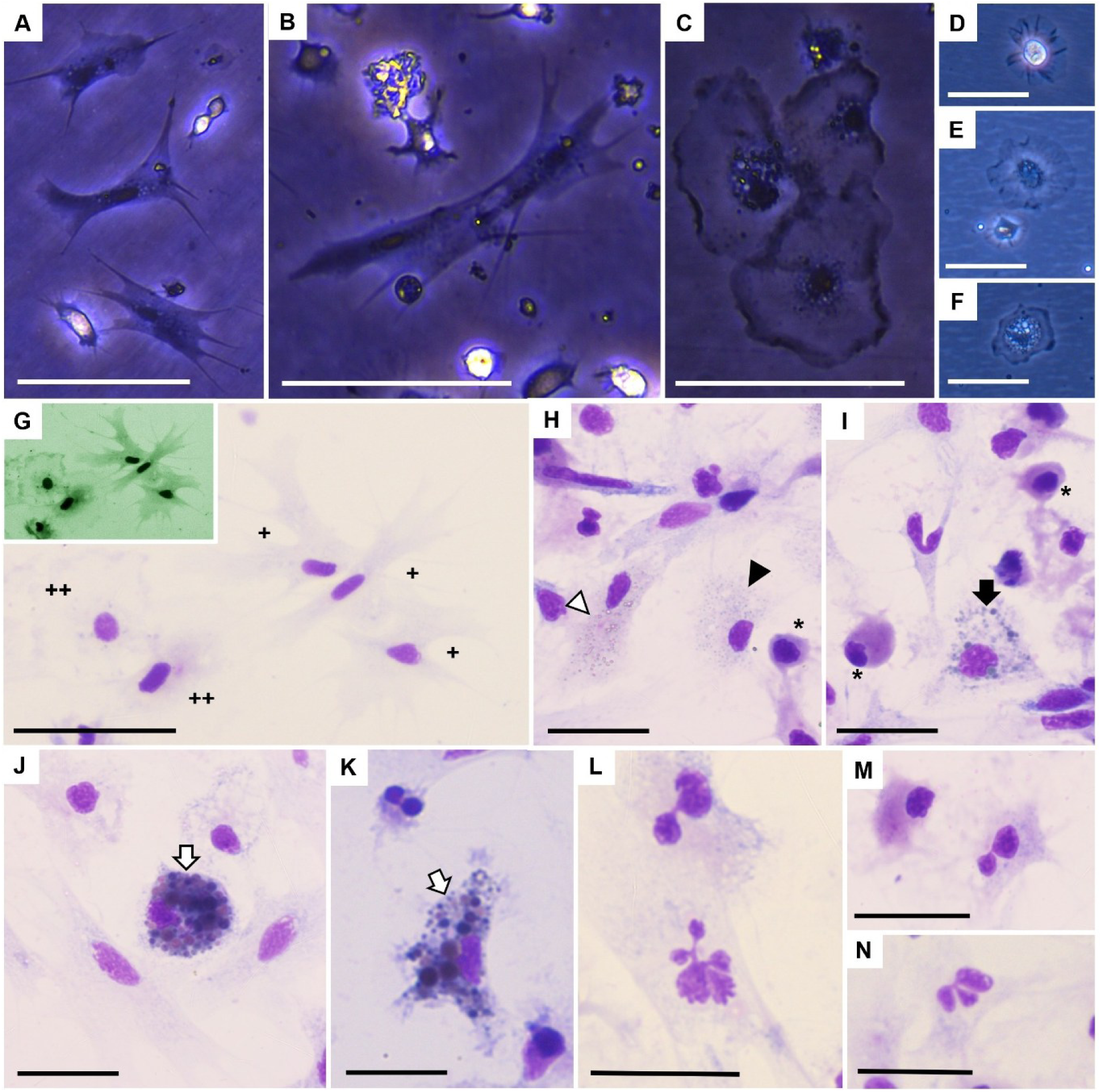
Morphological diversity of *Ostrea edulis* haemocytes. **(A-B)** Hyalinocyte-like haemocytes from fresh haemolymph, characterized by dark cytoplasm, fibroblast-like adherence, and fine filopodia; cells often overlapped upon contact. **(C)** Granulocyte-like haemocytes from fresh haemolymph, displaying rounded morphology, distinct nuclei, and variable cytoplasmic granulation; cells typically aggregated margin-to-margin. **(D-F)** Granulocyte-like haemocytes shed from explant cultures: initial adhesion (D), limited granulation (E), or extensive granulation (F). **(G-K)** Hemacolor-stained haemolymph preparations showing cytoplasmic heterogeneity: hyalinocyte-like haemocytes without granulation (G, single plus), degranulated granulocytes without granulation (G, double plus; inset: thallium LUT highlighting cell margins), blast-like cells without granulation (H-I, asterisk) refractive eosinophilic inclusions (H, open arrowhead), fine basophilic granules (H, closed arrowhead), moderate basophilic granules (I, closed arrow), and coarse basophilic granules (J-K, open arrows). **(L-N)** Hemacolor-stained haemolymph preparations revealed heterogeneity in polymorphonuclear cells, including bilobed (L-M), trilobed (N) and multilobed (L) forms. Scale bars: (A-G) = 50 µm; (H-N) = 20 µm.

In culture, granular haemocytes demonstrated marked plasticity. During degranulation, granulocytes lost their characteristic intracellular granules within 12 minutes, adopting an agranulocyte-like phenotype, while agranular granulocyte haemocytes could acquire intracellular granules within less than a minute (S4 Video). Additional modes of granulation were also observed, including membrane-associated uptake events consistent with endocytosis (S5 Video). Haemocytes were also frequently motile, transitioning between fully adherent and suspended states. The mean diameter of adherent haemocytes at their widest point was 34.96 ± 6.19 µm (range: 21.36-54.78 µm), encompassing both granular and degranulated forms. Diameters of suspended cells could not be reliably measured due to the presence of mixed cell types and transitional states.

### Cell diversity and biomineralization across cultured tissues

Fibroblast-like cells were observed across all explant cultures and provisionally grouped into five morphological categories (Types I-V), as their precise identities could not be confirmed. The occurrence of these categories varied, with some restricted to specific explant types. Type I cells were exclusive to heart-derived cultures (Fig 5A). They were often bordered by smaller cells with thin dendrite-like projections, designated Type II (Fig 5B). Type I cells maintained a distinct morphology but often formed loose associations with surrounding cells and occasionally developed dense patches with overlapping connections (Fig 5C). Type II cells were smaller, weakly adherent, and highly sensitive to medium changes. They frequently detached from the substrate, establishing only transient contact with haemocytes or other fibroblast-like cells (Fig 5C; S4 Video). Type III cells, abundant in gill-, mantle-, and heart-derived cultures, were typically observed as stand-alone cells with broad, flowing projections (Fig 5D). Type IV cells, restricted to heart cultures, resembled mature cardiomyocytes. They were smooth, elongated, and occasionally aligned in stacked arrangements with adjacent cells, although they did not contract (Fig 5E-F). Type V cells, readily adherent and therefore more amenable to Hemacolor staining, had lightly stained cytoplasm with multiple fine processes that interconnected to form multicellular aggregates (Fig 5G-I). These aggregates frequently expanded into adherent, scaffold-like networks (Fig 5J), which occasionally became infiltrated by additional cell types in heart cultures (Fig 5K). In heart cultures, aggregates further developed into large, interconnected clusters that exhibited spontaneous contractile activity (Fig 5L) or large monolayer-like networks (Fig 5M). Although the precise cellular composition of these larger networks could not be resolved, their organization and behaviour were distinct from smaller fibroblast-like networks, which remained non-contractile.

**Fig 5.**
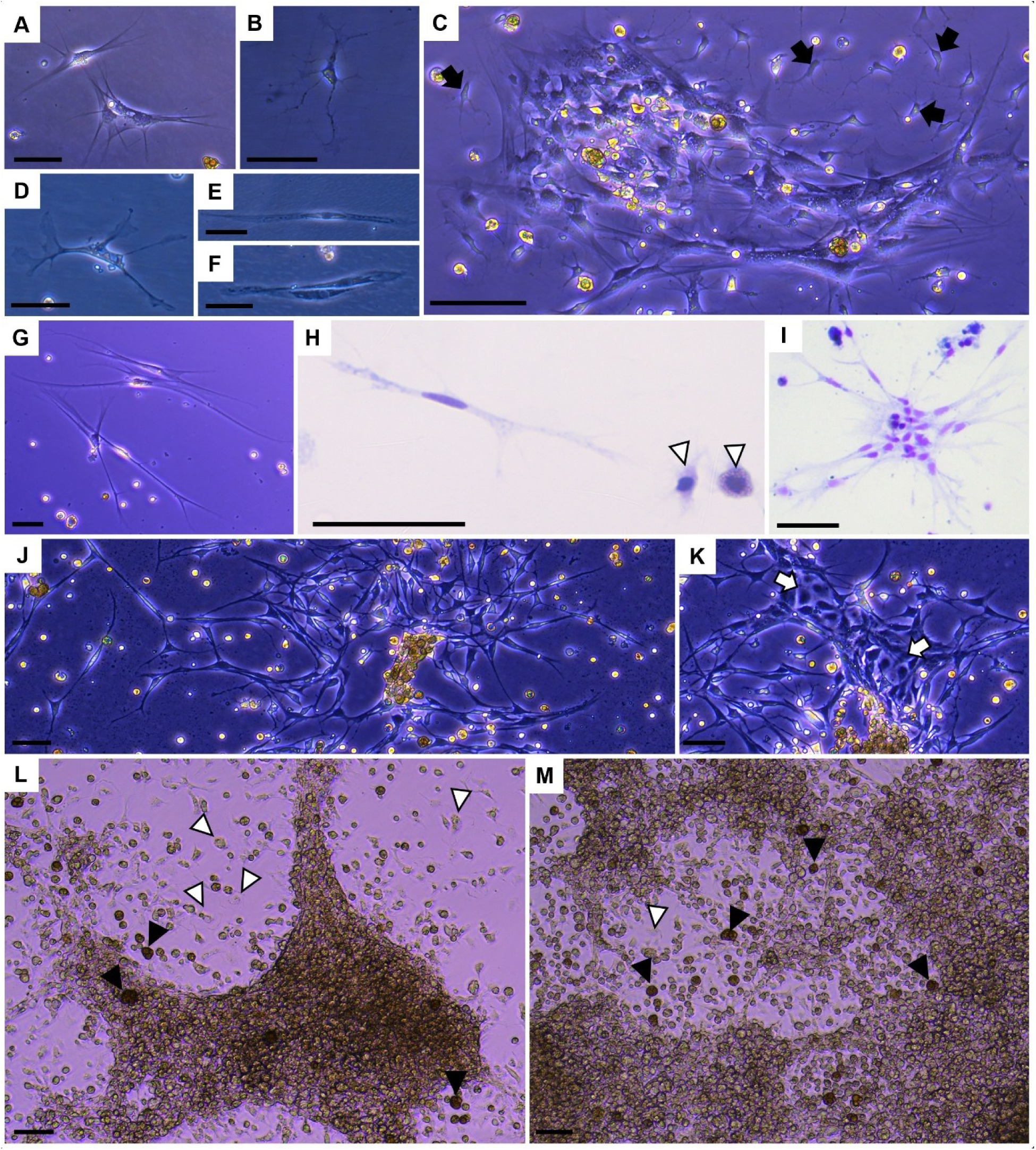
Fibroblast-like cell diversity observed in *Ostrea edulis* explant cultures. **(A)** Type I cells, exclusive to heart-derived cultures. **(B)** Type II cells, smaller with thin dendrite-like projections; these cells showed weak adherence and frequently detached. **(C)** Type I cells forming dense overlapping patches, with smaller Type II cells at the periphery (closed arrows). **(D)** Type III cells, abundant across gill-, mantle-, and heart-derived cultures, typically stand-alone with broad, flowing projections. **(E-F)** Type IV cells, restricted to heart cultures, resembling cardiomyocyte-like cells with smooth, elongated morphology and alignment into stacked arrangements. **(G-J)** Type V cells, readily adherent and therefore more amenable to staining, showing lightly stained cytoplasm and multiple fine processes that interconnected to form multicellular aggregates and network-like structures. Haemocytes indicated by open arrowhead **(K)** Type V fibroblast-like networks infiltrated by additional cell types in heart cultures (open arrow). **(L-M**) Large, contractile multicellular clusters arising in heart cultures. Additional cell types were also present, including haemocytes (open arrowheads) and auricle-associated granular cells (closed arrowheads). Scale: (A-M) = 50 µm.

In both mantle- and gill-derived cultures, individual ciliated cells were present in the medium as small, rounded cells bearing multiple motile cilia localized to one pole (Fig 6A-B). Following decontamination, cilia were initially lost in all explants but consistently regrew within two weeks of culture. In mantle explants, an outer necrotic layer was formed during decontamination, resulting in loss of the original ciliated mantle edge (Fig 6C-E). Rounded tissue masses subsequently developed, and new ciliated lobes emerged at one pole of the tissue explant (Fig 6D-E). These regenerated lobes did not replicate the full extension capacity or multi-layered lobed organisation of the original mantle edge but consistently re-established cilia along mantle margins. Weak and irregular contractions of mantle explants were sometimes observed but were unresponsive to external stimuli (e.g. media change), contrasting with the consistent, stimulus-responsive contractions of cultured heart tissue. Gill explants retained structural integrity for shorter periods, typically up to two months, although cells shed into the media remained viable beyond this point, up to a month after the parent tissue disassociated. Shed populations comprised mainly haemocytes, ciliated cells, and fibroblast-like cells resembling Type III. In addition to ciliated cells, mucous production was frequently observed during the early stages of both mantle- and gill-derived cultures and corresponded with the presence of mucocytes in the explanted tissues (Fig 6F-H). Cytological examination of cells shed into the medium from mantle and gill explants revealed a distinct eosinophilic cell type with foamy cytoplasm, which may represent mucocytes; as they were only present from cells shed into media of mantle and gill explants (Fig 6G-I). However, its identity could not be fully separated in live cell imaging from other suspension cells with a round phenotype (Fig 6K). An additional cell type, often found adjacent to the mucocytes in gill tissue and occasionally found within VM tissue and on the margin of heart ventricle tissue, exhibited a distinct eosinophilic phenotype. It did not stain positive for mucin and was very rarely found shed into the media (Fig 6H, 6J).

**Fig 6.**
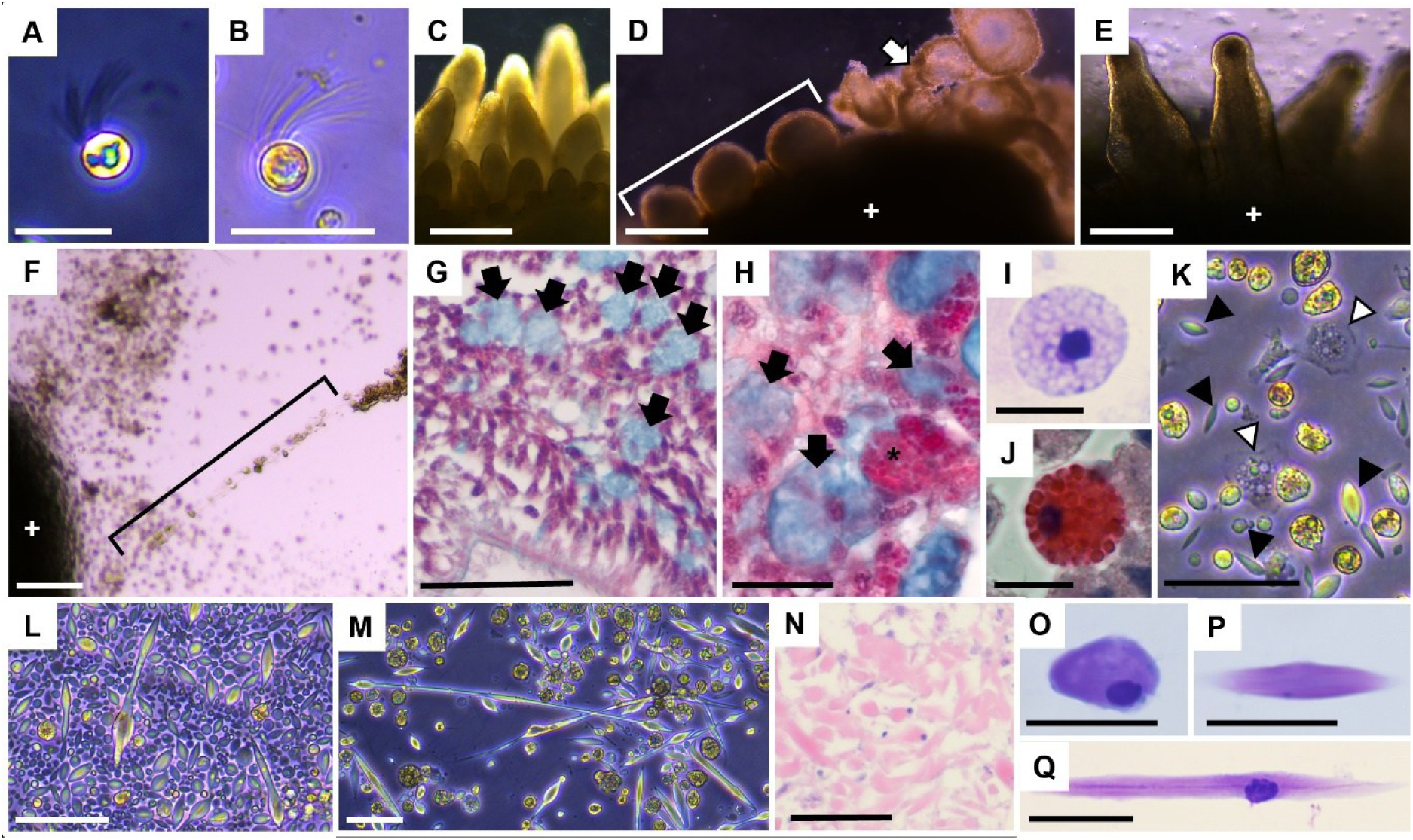
Cellular and Tissue Features of *Ostrea edulis* Mantle and Gill Explant Cultures. **(A-B)** Ciliated cells shed into culture medium. **(C)** Mantle lobes *in vivo*. **(D-E)** Mantle lobe regeneration in culture; regrowing lobes (bracket), outer dead layer shed into medium (open arrow), explant (plus). **(F)** Mucus in culture (bracket), explant; gill (plus). **(G)** Mantle tissue stained with Masson’s trichrome modified with Alcian blue; mucocytes (closed arrow). **(H)** Gill tissue stained with Masson’s trichrome modified with Alcian blue; mucocytes (closed arrow) and epithelial-associated eosinophilic cell (asterisk). **(I)** Hemacolor staining of a mantle and gill-derived cell shed into culture medium; likely a mucocyte. **(J)** Hemacolor staining of an epithelial-associated eosinophilic cell shed from gill explant culture medium. **(K)** Unidentified round cell phenotypes. Granulocytes (open arrowhead) and myocytes (closed arrowhead). **(L)** Smooth muscle cells (myocytes) in early culture, showing elliptical structures formed by proliferating myocytes **(M)** Elongated myocytes shed into medium, arising from rounded cells shown in (L). **(N)** Myocytes in mantle tissue from cultured explant, H&E stain. **(O-P)** Shed myocytes from culture, Hemacolor stain. Scale bars: (A, H, O, P) = 20 µm; (B, G, K, L, M, N, Q) = 50 µm; (C-E) = 500 µm; (F) = 100 µm; (I, J) = 10 µm.

Mantle explants gave rise to a distinct population of smooth muscle cells, myocytes (Fig 6L-M). In early culture, these appeared as small, round cell bodies measuring 3.65 ± 1.04 µm in diameter (range: 2.04-6.95 µm; S2 Table) that expanded outward in opposing directions, producing elliptical structures measuring 73.82 ± 93.13 µm in length (range: 7.79-576.42 µm; S2 Table) (Fig 6L-M). Elongated, elliptical myocytes were also shed directly into the medium from the explant, where Hemacolor and H&E staining confirmed strong similarity to tissue-resident myocytes (Fig 6N-Q). Shed myocytes did not exhibit contractile activity and progressively lost structural definition before dying; however, turnover was evident, as new myocytes continued to emerge during culture. Often, myocytes shed into the media exhibited a shrinkage of nuclei (Fig 6P).

In VM explant cultures, the predominant cells shed into the medium were spherical, non-adherent adipocytes, measuring 27.07 ± 4.71 µm in diameter (range: 17.45-40.04 µm; S2 Table). These cells displayed polarized, refractile cytoplasm and sometimes occurred in closely apposed pairs (Fig 7A-B). Their identity was confirmed by Histogel embedding of shed cells (corresponding to those shown in Fig 7A; Fig 7C) and comparison with the resident adipocyte population in the fresh and explanted tissue (Fig 7D). Adipocytes maintained a large, rounded, non-adherent morphology for up to seven months in culture. Beyond this period, some cells adopted an elongated, adherent phenotype, anchoring the explant to the tissue culture plate (Fig 7E-G). The appearance of these elongated cells was associated with culture instability and an increased risk of complete tissue dissociation. In addition to adipocytes, the second most prevalent cell types shed from VM explants were haemocytes, present as both granular (Fig 7H) and agranular or degranulated granulocytes (Fig 7I), as well as fibroblast Type II cells (Fig 7I). Although the cells shed into the media largely represented haemocyte and adipocyte cells, several VM explants also shed a large, opaque cell population whose identity could not be positively confirmed with live cell imaging or histology (Fig 7J-K).

**Fig 7.**
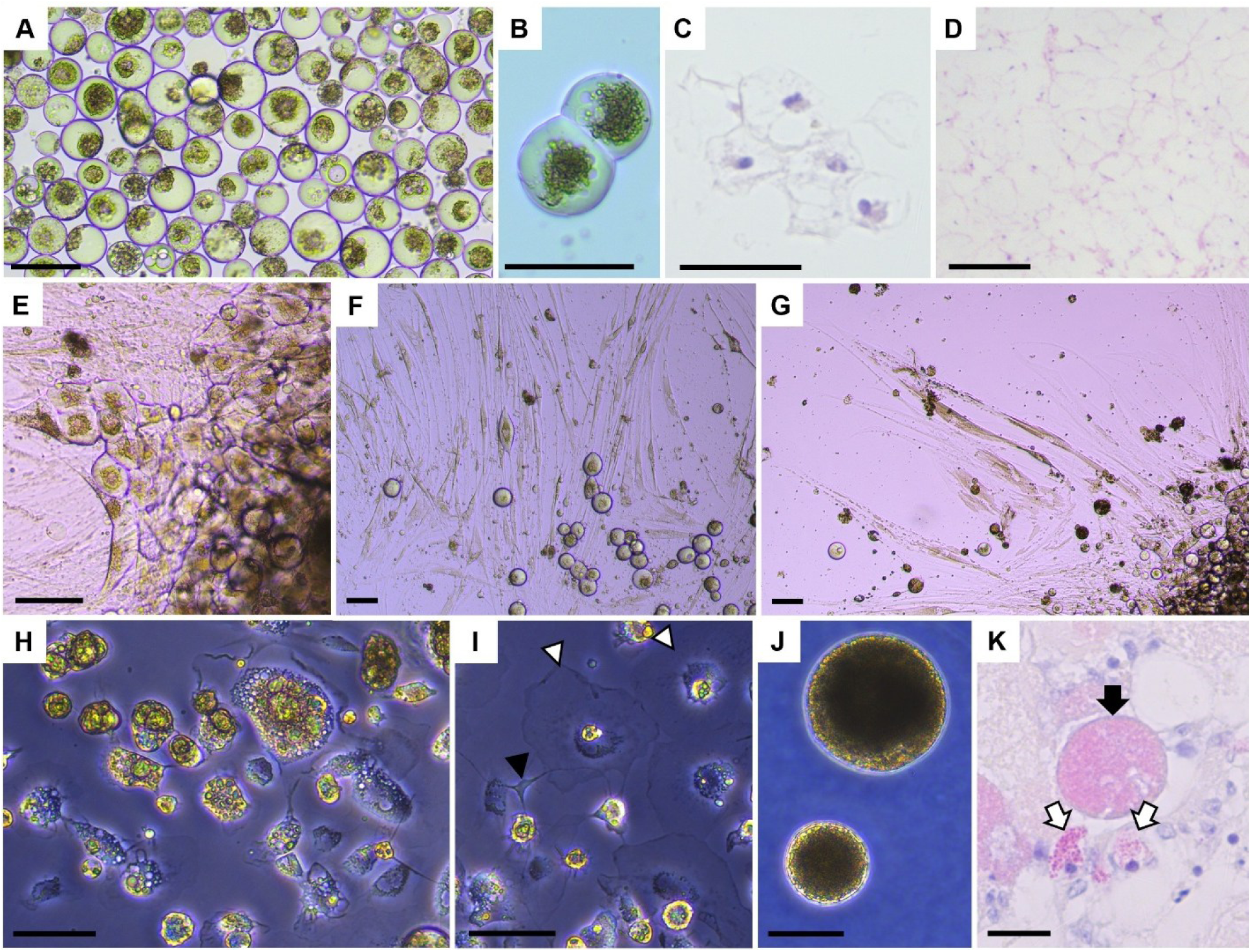
Visceral Mass (VM) explant culture features in *Ostrea edulis*. **(A)** Non-adherent adipocyte cells. **(B)** Adipocyte cells with polarized cytoplasm. **(C)** Adipocyte cells shed into culture medium, embedded in Histogel, H&E (corresponding to cells in A). **(D)** Adipocyte tissue within cultured explant, H&E. **(E)** Adherent explant showing transitional adipocyte morphology. **(F-G)** Monolayer of intermediate adipocyte forms and elongated cells. **(H)** Granular haemocytes. **(I)** Degranulated granular haemocytes (open arrowhead) and fibroblast type II (closed arrow). **(J)** Large, undefined cell type specific to prolonged visceral mass cultures. **(K)** H&E stained 3-month cultured explant highlighting large, undefined cell type (closed arrow) and eosinophilic cells (open arrow). Scale bars: (A-C, E-J) = 50 µm; (D) = 100 µm; (K) = 20 µm.

Hard structured mineralized material formed in multiple explant cultures, regardless of tissue origin (Fig 8A-F). These shell-like fragments were often surrounded by clusters of adherent cells. Morphologies varied with tissue source: heart and mantle produced highly refractive, nacre-like material (Fig 8A-C); heart and visceral mass also generated dense, opaque fragments (Fig 8A-B, 8D); and gill uniquely formed fragments with a gradient, appearing heavily mineralized at one end and increasingly transparent toward the other (Fig 8E-F). The fragments remained stable throughout culture and were unaffected by media changes or freshwater exposure. No mineralized or precipitate-like material was observed in control wells, indicating that formation required the presence of explanted tissue.

**Fig 8.**
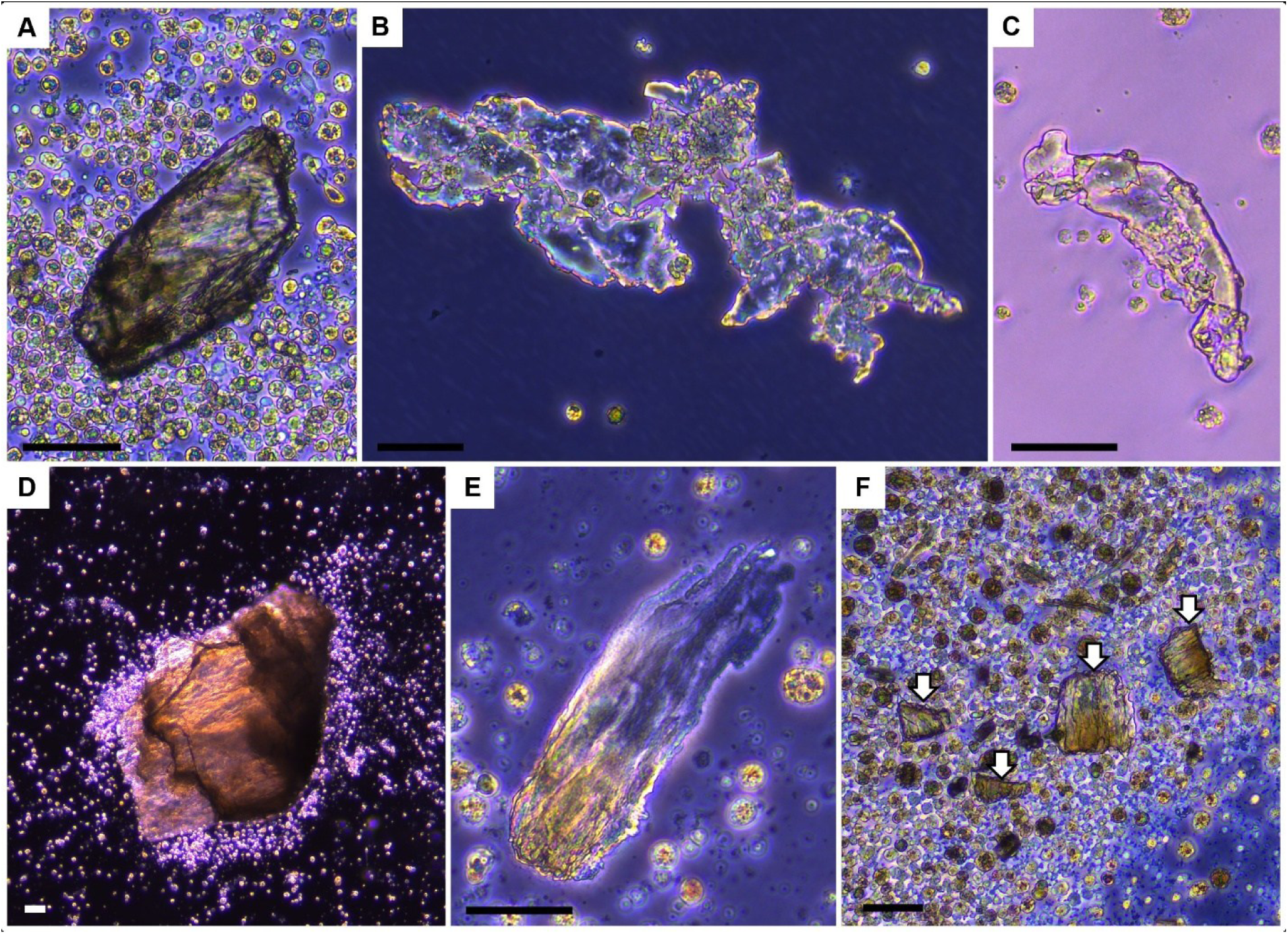
*De novo* Formation of Shell-like Mineralised Structures in *Ostrea edulis* Tissue Cultures. **(A-B)** Originating from heart explants. **(C)** Originating from mantle explant. (D) Originating from VM explant. **(E-F)** Originating from gill explants (closed arrow). Scale bar: (A-F) = 50 µm.

### Crosscutting culture challenges

Contamination was most common during the first 24 h of culture but became minimal once explants became established. Bacterial contamination was the most frequent, followed by small, unidentified ciliates, both occurring only during the first few days of culture. Long-term maintenance was generally reliable among explants that remained viable beyond this initial period. By 24 h post-processing, viability could usually be determined: explants that remained free of contamination at this stage typically survived and developed further.

Among successfully established cultures, approximately 75% could be maintained to the maximum durations observed: mantle up to 4 months, gill up to 2 months, VM up to 8.5 months, and contracting hearts up to 9 months. Many cultures were terminated earlier for histological or imaging analyses, so these values reflect achievable longevity rather than the lifespan of every explant.

Contamination rates varied by tissue. It occurred in approximately 30-40% of gill cultures, 20-30% of mantle cultures, 15% in VM, and <5% in heart explants. All adductor muscle cultures failed due to persistent bacterial contamination within the first 24 hours. These percentages represent estimates from repeated culture attempts.

In addition to these technical considerations, all explant cultures consistently released abundant small non-adherent cells into the medium. These were morphologically distinct from pigmented auricle-associated granule cells and adipocytes. At least a subset was consistent with haemocytes in their non-adherent form; however, without definitive markers, haemocytes could not be reliably distinguished from other non-adherent cell types, and vice versa.

## Discussion

The establishment of long-term oyster tissue cultures represents a major step toward understanding molluscan cell biology. By sustaining diverse tissue types and cell populations *in vitro*, these systems enable detailed study of cellular adaptation, plasticity, and function, opening new avenues for physiological, pathological, and aquaculture research. Across explant types, heart and VM exhibited the greatest longevity and stability, with contractile hearts maintained for up to nine months and VM cultures for eight and a half months. Mantle explants remained viable for up to four months, while gill explants retained tissue integrity for up to two months. Cell populations shed into the media reflected their tissue origins: hearts produced cardiomyocyte-like cells, fibroblast-like subtypes, and haemocytes; VM cultures yielded adipocytes and haemocytes; mantle explants generated myocytes, fibroblast-like cells, and haemocytes; and gill cultures primarily released haemocytes and fibroblast-like cells. Collectively, these results demonstrate that multiple oyster tissues and cell types can be maintained long-term under common culture conditions, yet each exhibits distinct patterns of survival, morphology, and behaviour. These extended culture periods further enabled direct observation of rhythmic contractility and novel ciliation in hearts, persistence of adipocytes in VM, biomineralization across tissues, and unexpected cellular plasticity, including haemocyte granule turnover and *de novo* microtissue formation. Together, these findings establish *O. edulis* explant cultures as a tractable and informative platform for both fundamental and applied research.

Importantly, the capacity of these explants to survive long-term is coupled with a measurable acclimatization process, revealing how oyster tissues physiologically adjust to *in vitro* conditions. A key finding of this work is the time-dependent acclimatization of explants. Isolated hearts required approximately 14 days in culture before heart rates stabilized, reflecting a period of adjustment likely driven by the stress of excision, adaptation to the artificial environment, and remodelling of cell-cell and cell-matrix interactions. Comparable lag phases have been reported in vertebrate tissue culture systems. For example, intact myocardial slices have been shown to retain contractile stability only after an initial equilibration period in culture ^31^. The presence of a defined adjustment window carries both biological and methodological significance: it demonstrates that oyster cardiac tissue can establish a new *in vitro* equilibrium, while highlighting that early measurements capture acute stress responses rather than steady-state physiology. Accordingly, functional assays are most reliable after this two-week stabilization period.

Although other tissues did not allow equivalent functional assessment, comparable adjustment processes were apparent. In mantle and gill explants, ciliated lobes and cilia, respectively, were initially lost but consistently regenerated within two weeks, suggesting a shared acclimatization timescale. The mantle provided a clear example of tissue-specific adaptation, displaying two hallmark behaviours that align with and extend current work on bivalve epithelia and muscle. The loss and subsequent restoration of lobes and ciliation are consistent with the intrinsic ciliogenic capacity and epithelial resilience of molluscan mantle cells ^32,33^. In parallel, the emergence of shed smooth-muscle cells that transiently proliferate yet lack sustained contractility echoes prior culture studies where myocyte function and differentiation depend on appropriate extracellular matrix and mechanical context ^33,34^]. Together, these findings suggest that mantle-derived cultures complement cardiac preparations as a model for molluscan muscle and offer a tractable system for investigating epithelial repair and ciliogenesis.

The *ex vivo* heart rates obtained in this study were substantially lower than those reported for *O. edulis* under natural conditions, where cardiac frequencies typically range between 10 and 40 bpm depending on temperature and activity ^35^. This discrepancy is expected, as the isolated heart is removed from the haemodynamic influences present in the intact organism, and may operate at a reduced basal level. Importantly, low heart rates and periods of cardiac suppression are not unprecedented in *O. edulis*. During valve closure or in response to unfavourable environmental conditions, oysters display pronounced bradycardia and even transient cardiac arrest ^35^. Therefore, although our cultured preparations show heart rates lower than *in vivo* values, these remain consistent with the physiological capacity of the species to down-regulate cardiac activity during dormancy or stress.

Collectively, these observations illustrate that successful acclimatization underlies long-term tissue stability in culture, providing the foundation for sustained physiological function. Once acclimatized, the cultured hearts not only stabilized but also retained contractile function over extended periods, demonstrating sustained physiological viability well beyond typical *in vitro* expectations. This longevity creates opportunities to observe long-term adaptations, remodelling, and emergent dynamics not accessible in short-lived culture systems.

Comparisons with other molluscs highlight both parallels and distinctions. In *Magallana (formerly Crassostrea) gigas*, excised whole hearts continued rhythmic contractions for up to eight weeks *in vitro*, although explants gradually rounded and preservation of auricle-ventricle architecture was uncertain ^26^. In *Magallana bilineata* (*formerly Crassostrea madrasensis*), small heart fragments produced migrating cells that aggregated into beating clusters surviving 25-34 days, but contractions weakened after four weeks ^24^. Such cluster-based systems differ from intact hearts, yet their behaviour parallels our observations of cells shed from whole-heart explants, which reorganized into *de novo* cardiac microtissues. Similar diversity is seen in the hard clam *Meretrix lusoria*, where fragmented heart tissues generated confluent epithelial-like monolayers after 30 days, occasionally exhibiting rhythmic contractions ^25^. In the scallop *Pecten maximus*, dissociated heart tissues produced cardiomyocyte-like cells that began spontaneous rhythmic contractions after approximately 20 days, with myotube-like structures appearing at confluence ^36^. These outcomes resemble the aggregated beating clusters of Balakrishnan et al. (2024) and parallel our own findings, in which aggregations of Type V fibroblasts cells formed scaffolding and exhibited rhythmic contractions when part of large formations while Type I fibroblast cells formed compact but non-contractile structures. Together, these studies illustrate the range of cellular organizations capable of sustaining contractile activity in molluscan heart cultures, from beating clusters to sheet-like monolayers, while complementing the unique longevity and structural preservation achieved in our *O. edulis* heart explants.

Auricle-derived granule cells and the spontaneous organization of microtissues were features unique to heart cultures. This finding is particularly noteworthy, as the cellular sources driving such growth and regeneration in molluscs remain unknown. Stem cell populations in these animals have not yet been clearly identified or localized, making our observations valuable for understanding potential mechanisms of tissue renewal *in vitro* ^37^.

Auricle-derived cultures were dominated by granule cells, which frequently contained multiple internal bodies within a single cytoplasmic boundary. Time-lapse recordings revealed that these multinuclear or multivesicular cells proliferated in close association with emerging microtissues. Live-cell imaging and histological analyses confirmed that granule cells were incorporated into the resulting, fully structured microtissues. Whether these cells mature to form the complete tissue or instead act as organizers initiating microtissue assembly remains uncertain. The resulting self-organized cardiac microtissues share several features with vertebrate organoids; however, unlike lumenized organoids, they retain central cells and are composed of diverse cell types, making them superficially similar to spheroids but distinct in cellular organization ^38^. Their emergence likely reflects plastic reorganization of differentiated cells, though a contribution from unrecognized progenitor cells cannot be excluded. The progenitor potential, identity, and functional role of the auricle-derived granule cells therefore remain unresolved. Their consistent presence in both freshly isolated and long-term cultured hearts suggests that they represent a native cardiac cell population, potentially participating in or responding to the tissue reorganization observed *in vitro*. Supporting this interpretation, Renault et al. (1995) reported that the “brown cells” of the auricle exhibit strong survival and growth capacity *in vitro*.

Terminology surrounding granular molluscan cells is inconstant, with overlapping descriptions appearing under various names, including “pigment cells,” “brown cells,” “pore cells,” and “morula-like cells” ^39,40^. Granule cells are well documented in the pericardial (Keber’s) glands of several bivalves, where vesicle-rich secretory and absorptive epithelia occur. The pericardial gland itself varies among species -reported as lying on the auricular surface, lining the pericardial cavity, or forming glandular out-pockets of the pericardial wall ^41^. In our histological analyses, no discrete glandular structure was detected, and to our knowledge, a species-specific description of a distinct pericardial gland *in O. edulis* has not been established. Comparable inclusion-rich cell types have also been described in other molluscs, notably the “serous” haemocytes, which share some functional features with vertebrate mast cells ^40^. Nevertheless, our histochemical analyses revealed no proteoglycan staining characteristic of mast cells, suggesting that the granular cells observed here are unlikely to be mast-cell analogues. Matozzo et al. (2025) described these “serous cells” as a rare haemocyte subtype (<1% of circulating cells) capable of degranulation and DNA trap formation for bacterial agglutination. The “brown cells” of *O. edulis* hearts reported by Renault et al. (1995) exhibited similar degranulation behaviour, as they appeared less pigmented and granular in culture than in freshly dissociated preparations. Consistent with this, diagnostic practices often favour ventricular over auricular imprints due to a higher abundance of serous-like cells, likely corresponding to the “brown cells” described by Renault et al. (1995) and auricle-derived granular cells described here ^42^. Whether these variously described populations represent a single, multifunctional cell type or reflect broader inconsistencies in molluscan cell classification remains unresolved. At present, no defined cell type can be unequivocally matched to the predominant auricular cells of *O. edulis*, beyond descriptive labels such as “granule,” “brown,” or “pigmented” cells.

Long-term maintenance of explants under *ex vivo* conditions was accompanied by structural remodelling, suggesting a degree of cellular plasticity or dedifferentiation. Histological analyses showed that the epicardial epithelium became thicker, with cells adopting a more cuboidal morphology compared to the flattened squamous cells of freshly dissected hearts. In addition, motile cilia were observed on the epicardial epithelium covering auricular regions, a feature absent in fresh preparations. Previous studies in mollusc gills typically describe ciliary loss, damage, or functional arrest, and in some cases epithelial metaplasia ^43,44^. To our knowledge, the appearance of motile cilia on an epithelial surface not normally ciliated has not been reported in molluscan tissues. In vertebrates, ciliogenesis can be re-initiated in epithelia where cilia are part of the normal developmental program ^45^. While the appearance of motile cilia on the epicardial epithelium may represent an instance of epithelial plasticity; it may also be a culture-related response to prolonged stagnant conditions, a delayed form of epithelial remodelling under stress, or metaplastic changes triggered by long-term maintenance. Furthermore, the progressive increase in collagen deposition within the *in vitro* heart culture indicates ongoing extracellular matrix remodelling. Further characterization of fibroblast activity, collagen organization, and matrix turnover would help clarify whether this trend represents the onset of maladaptive remodelling. Another feature of long-term preparations was the appearance of hyaline globules along the auricular perimeter, often at ventricular margins and adjacent to remodelled epithelia. These structures were absent in freshly dissected hearts, suggesting that they arise, or become more prominent, under culture conditions. Their proximity to ciliated regions raises the possibility of a link with localized remodelling or secretory activity, although their cellular origin and physiological role remain uncertain. Alternatively, they may reflect culture-related changes in membrane permeability, leading to altered secretion, uptake, or accumulation of material. Superficially similar structures have been described in vertebrates, including Mallory-Denk bodies in hepatocytes and eosinophilic globules in tissues of cetaceans ^46,47^. Both of these vertebrate examples, however, are intracellular inclusions, whereas the oyster hyaline globules appear extracellular, and any resemblance should therefore be considered analogy rather than homology. While the nature of these globules remains uncertain, they form part of a broader pattern of tissue remodelling and cellular activity observed in long-term cultures.

In parallel, the explants gave rise to multiple cell types, including fibroblast-like cells, adipocytes, myocytes, haemocytes, and putative cardiomyocytes, whose organization and behaviour could be observed more clearly than in short-lived preparations. Shed cells could also be traced back to their tissue of origin; for example, adipocytes were characteristic of VM explants, while myocytes were a defining feature of mantle explants. Establishing such correlations between cultured cells and their source tissues may help refine existing terminology, particularly for mantle-derived myocytes, which in culture range from elliptical to fusiform morphologies and have often been described collectively under the broad term “spindle cells” ^26,48,49^.

Among the diverse populations shed into the media, haemocytes were particularly notable for their dynamic behaviour in culture. While haemocytes of oysters and other bivalves are well studied *in vivo,* their activity *in vitro* revealed an unexpected degree of phenotypic plasticity. Granulocytes rapidly lost their characteristic inclusions within minutes, producing degranulated forms that resembled agranulocytes. Some of these cells later reacquired granules, suggesting a reversible transition between granular and agranular states. True agranulocytes (hyalinocytes), however, were not stably maintained under our culture conditions, indicating that they may require distinct environmental or signalling cues for survival. This finding refines current models of haemocyte plasticity by demonstrating that degranulation can occur rapidly and reversibly *in vitro*, whereas certain haemocyte subtypes may be selectively lost in culture. Such dynamic transitions may also occur *in vivo* but remain undetected by static or population-level analyses ^8,50,51^.

Haemolymph cytology confirmed two principal haemocyte classes: hyalinocytes and granulocytes. The granulocyte population displayed marked heterogeneity, encompassing cells across a continuum from agranular to highly granular morphologies. Within this spectrum, several subtypes were distinguished based on staining characteristics and cytoplasmic inclusions, including agranular granulocytes, cells with refractive eosinophilic inclusions, and those containing fine, moderate, or coarse basophilic granules. The presence of these diverse cytological profiles suggests that granulocytes may comprise multiple discrete cell types or represent different physiological states within a single, plastic lineage. All granulocyte subtypes were observed in culture, whereas true hyalinocytes were absent, supporting the idea that they do not persist under these culture conditions. Collectively, these findings have important implications for *in vitro* functional and disease modelling studies, underscoring the need to account for haemocyte plasticity and the selective loss of specific subtypes when interpreting immune cell behaviour.

More broadly, these findings accentuate the dynamic and multifunctional nature of oyster cells in culture, revealing degrees of plasticity that extend beyond immune processes. Such cellular versatility was also evident in unexpected contexts, including biomineralization, where mineralized material developed across multiple explant types. In bivalves, shell deposition typically occurs at the periostracal groove, where specialized mantle epithelia coordinate secretion and calcium carbonate crystallization ^52^. In culture, however, new mineral deposits appeared irrespective of tissue origin. Given the extensive washing procedures and the direct observation of their formation during culture, these structures cannot be explained by residual shell fragments. Similar mineralization has been reported in mantle-derived cultures, where cells secrete calcium carbonate or organic matrix components ^53^. Although haemocytes are generally associated with immune functions, they are also known to contribute to shell repair and calcium transport ^8,54,55^. Recent work has identified stem-like “skeleton-embedding cells” within molluscan shells that exhibit both hematopoietic and biomineralizing potential ^56^. Although our data do not directly confirm the presence or activity of specialized biomineralizing cells, the observation of mineral deposition across multiple explant types suggests a broader capacity for biomineralization within molluscan tissues. This pattern is consistent with previous evidence that multiple cell types in molluscs can exhibit mineralizing activity under certain conditions ^57^. These findings raise the possibility that a latent mineralizing potential may be an intrinsic property of several oyster cell populations, extending beyond the mantle where shell formation is classically localized. Further studies will be needed to determine whether comparable progenitor or stem-like populations contribute to this phenomenon under culture conditions.

Collectively, these findings demonstrate that cells shed in oyster cultures are not passive survivors but dynamic participants in tissue maintenance, remodelling, and even *de novo* structure formation. The plasticity of haemocytes, the diversity of fibroblast-like cells, the persistence of adipocytes, the regenerative capacity of mantle tissues, and the mineralizing activity observed across explant types all point to a broad cellular repertoire that extends beyond traditional expectations for molluscan primary cultures. At the same time, the identities and functional roles of many of these cell types remain unresolved. Long-term culture systems therefore provide a platform to disentangle the contributions of different populations to immunity, metabolism, regeneration, and biomineralization.

This diversity, however, also highlights the interpretive limits of morphology-based classification in primary cultures. In mantle and gill explants, visible mucous material was observed, and Alcian Blue staining confirmed the presence of mucocytes in the source tissues. Secretion-like material occasionally emanated from explants *in vitro*, but whether mucocytes themselves were shed into the medium could not be determined. It remains possible that some of the rounded cells released from explants were mucocytes, illustrating the broader difficulty of interpreting cell identity in these cultures. The frequent use of “rounded” as a descriptor exemplifies this challenge ^26,58,59^. Rounded morphology may arise from multiple cell types, yet in the absence of lineage-specific markers such diversity collapses into a single ambiguous category, thereby masking important functional differences. Improved molecular tools will be essential to resolve the identities of cell populations and move beyond generalized morphological labels.

In addition to interpretive challenges, practical limitations remain in establishing and maintaining certain tissue types under these conditions. Although the culture conditions supported several tissue types, we were unable to establish cultures from adductor muscle. This difficulty appears to be specific to the adductor rather than muscle-derived cells in general and may be related to the compact structure of this tissue. Further work will be needed to determine whether alternative culture approaches could improve outcomes. Additionally, it remains uncertain whether similar outcomes can be replicated across other molluscs. Nonetheless, such challenges present opportunities: extending culture methods across molluscan taxa may improve outcomes for difficult tissues and reveal differences in cell types and organization. In this way, comparative approaches can both expand the technical toolkit for molluscan cell culture and deepen our broader understanding of invertebrate cell biology.

Developing long-term cultures of *O. edulis* tissues brings molluscs closer to other invertebrate model systems and establishes a practical foundation for applied and fundamental research, paving the way for future advances in aquaculture, cell biology, and biotechnology.

## Acknowledgements

The authors gratefully acknowledge Dr. Anna Raper of the Roslin Institute Bioimaging and Flow Cytometry Facility for her expertise and technical assistance. We thank John P. Bignell and Dr. David Bass of the Centre for Environment, Fisheries and Aquaculture Science (CEFAS), and Victoria A. Sleight of the University of Aberdeen for their constructive scientific discussions. We are also grateful to the Royal (Dick) School of Veterinary Studies Large Animal Research and Imaging Facility (LARIF) for access to equipment and the Histopathology Laboratory, Diagnostic Biopsy Service, Royal (Dick) School of Veterinary Studies, University of Edinburgh, for processing the histological samples. For the purpose of open access, the author has applied a CC-BY public copyright licence to any Author Accepted Manuscript version arising from this submission.

## Funding Statement

The authors acknowledge funding from the Biotechnology and Biological Sciences Research Council (BBSRC); including an EASTBIO DTP Studentship [BB/T00875X/1] to KRS, a BBSRC Discovery Fellowship [BB/X009807/1] to PCS, and an Institute Strategic Programme grant [BBS/E/RL/230001C].

## Competing Interests

The authors declare no completing interests.

## Author Contributions

[TPB;TR;KRS] Conceptualization, [KRS] Data Curation, [KRS] Formal Analysis, [TPB; TR; LJM; PCS] Funding Acquisition, [KRS; SM] Investigation, [TPB; TR; LJM; PCS; SM; KRS] Methodology, [TPB] Project Administration, [TPB; KS] Resources, [TPB; TR; LJM; PCS] Supervision, [KRS] Validation, Visualisation, [KRS] Writing – Original Draft Preparation, [TPB; TR; LJM; PCS; SM; KRS] Writing – Review & Editing.

## Abbreviations

AE: auricular epicardium
AM: adductor muscle
ASW: artificial seawater
AU: auricle
Bge: Biomphalaria glabrata embryonic
BP: biopsy punch
CfT: Chlamys farreri trochophore
GI: gill
H&E: haematoxylin and eosin
HE: heart
MA: mantle
OE: Ostrea edulis
VE: ventricle
VM: visceral mass.

## Supplementary Information

### Videos

**S1 Video. *Ex vivo* culture *Ostrea edulis* heart auricle showing novel ciliation and fluid movement.** Video recording of the auricular region from an *ex vivo* cultured *Ostrea edulis* heart. Cilia are visible along the tissue margin, generating directional currents that move cells and debris at the tissue edge. The observation reveals previously undescribed ciliation associated with the auricle under prolonged culture conditions.

**S2 Video. Contracting microtissues derived from *Ostrea edulis* heart cultures.** *De novo* microtissues originating from primary *Ostrea edulis* heart explant cultures show spontaneous contractile activity. **(A)** Example of a small contracting microtissue (scale bar: 50 µm). **(B)** Example of a large contracting microtissue maintained in long-term culture (scale bar: 250 µm). **(C)** Example of an adherent contracting microtissue maintained in long-term culture (scale bar: 100 µm). **(D)** Microtissue exhibiting spontaneous cilia formation (scale bar: 100 µm).

**S3 Video. Early microtissue formation.** Time-lapse video composed of images captured automatically every 15 minutes over 48 hours and stitched together in ImageJ, showing microtissue growth in culture. The inset highlights an auricle-derived granule cell containing multiple internal bodies undergoing division.

**S4 Video. Dynamic granulation of cultured haemocytes.** Time-lapse video (2 h 15 min total duration) composed of images captured manually every 30 seconds and stitched together in ImageJ. In the first 5 seconds of playback, (i) shows a granulated haemocyte undergoing degranulation and transforming into a cell with an agranular phenotype within approximately 15 minutes of real time. The asterisk (*) highlights a Type II fibroblast with dendrite-like projections interacting with haemocyte membranes. In the final 5 seconds of playback, (ii) shows an agranular haemocyte rapidly acquiring granules, representing approximately 1 minute of real time. Scale bar: 50 µm.

**S5 Video. Haemocyte granulation through endocytosis.** Time-lapse video composed of images captured manually every 30 seconds over a 40-minute period and stitched together in ImageJ. The sequence shows active membrane endocytosis and progressive granule accumulation within a cultured haemocyte. Scale bar: 100 µm

### Tables

**S1 Table. Heart rate measurements before and after media replacement.** Raw dataset showing individual heart rate measurements recorded before and after media changes. Five cultured hearts were recorded once before and once after each media replacement. Contractions were counted over a 2-minute period and divided by two to calculate beats per minute (bpm). Measurements were collected weekly, corresponding to days 0, 7, 14, 21, 28, and 35 in culture.

**S2 Table. Raw cell and granule measurement dataset from *Ostrea edulis* tissues and primary cultures.** Dataset compiling quantitative measurements of cell and particle diameters from Ostrea edulis tissues and in vitro cultures. Data include granules, granule aggregates, hyaline globules, auricle-derived granule cells, adherent haemocytes, mantle myocytes (early- and late-stage growth), and adipocytes.

## Data availability statement

All relevant data are within the paper and its Supplementary Information files.

## Notes

### Competing Interest Statement

The authors have declared no competing interest.

## References

1. MolluscaBase. Published online 2024. Accessed September 12, 2025. https://www.molluscabase.org

2. Vaughn CC, Hoellein TJ. Bivalve Impacts in Freshwater and Marine Ecosystems. Annu Rev Ecol Evol Syst. 2018;49(1):183–208. doi:10.1146/annurev-ecolsys-110617-062703

3. Jung DY, Kim HG, Kim TL, et al. Blue carbon potential of oyster: Review of global oyster production and annual growth rate. Regional Studies in Marine Science. 2025;86:104195. doi:10.1016/j.rsma.2025.104195

4. Van Der Schatte Olivier A, Jones L, Vay LL, Christie M, Wilson J, Malham SK. A global review of the ecosystem services provided by bivalve aquaculture. Reviews in Aquaculture. 2020;12(1):3–25. doi:10.1111/raq.12301

5. Davison A, Neiman M. Mobilizing molluscan models and genomes in biology. Phil Trans R Soc B. 2021;376(1825):20200163. doi:10.1098/rstb.2020.0163

6. Balakrishnan S, Singh ISB, Puthumana J. Status in molluscan cell line development in last one decade (2010–2020): impediments and way forward. Cytotechnology. 2022;74(4):433–457. doi:10.1007/s10616-022-00539-x

7. Yoshino TP, Bickham U, Bayne CJ. Molluscan cells in culture: primary cell cultures and cell lines. Can J Zool. 2013;91(6):391–404. doi:10.1139/cjz-2012-0258

8. De La Ballina NR, Maresca F, Cao A, Villalba A. Bivalve Haemocyte Subpopulations: A Review. Front Immunol. 2022;13:826255. doi:10.3389/fimmu.2022.826255

9. Ladhar-Chaabouni R, Hamza-Chaffai A. The cell cultures and the use of haemocytes from marine molluscs for ecotoxicology assessment. Cytotechnology. 2016;68(5):1669–1685. doi:10.1007/s10616-015-9932-3

10. Metzger MJ, Reinisch C, Sherry J, Goff SP. Horizontal Transmission of Clonal Cancer Cells Causes Leukemia in Soft-Shell Clams. Cell. 2015;161(2):255–263. doi:10.1016/j.cell.2015.02.042

11. Rinkevich B, Pomponi SA. Advancing marine invertebrate cell line research: four key knowledge gaps. In Vitro CellDevBiol-Animal. 2025;61(5):493–505. doi:10.1007/s11626-025-01029-y

12. Ashton-Alcox KA, Ford SE. Variability in molluscan hemocytes: a flow cytometric study. Tissue and Cell. 1998;30(2):195–204. doi:10.1016/S0040-8166(98)80068-2

13. Lu J, Shi Y, Yao T, Bai C, Jiang J, Ye L. Gender Differences in Hemocyte Immune Parameters of Hong Kong Oyster Crassostrea hongkongensis During Immune Stress. Front Immunol. 2021;12:659469. doi:10.3389/fimmu.2021.659469

14. Castillo MG, Humphries JE, Mourão MM, Marquez J, Gonzalez A, Montelongo CE. Biomphalaria glabrata immunity: Post-genome advances. Developmental & Comparative Immunology. 2020;104:103557. doi:10.1016/j.dci.2019.103557

15. Hansen EL. A Cell Line from Embryos of Biomphalaria glabrata (Pulmonata): Establishment and Characteristics. In: Invertebrate Tissue Culture. Elsevier; 1976:75–99. doi:10.1016/B978-0-12-470270-7.50011-2

16. Qin Z, Ji A, Yan M, et al. Establishment of the first marine mollusk cell line from scallop (Chlamys farreri) trochophore. Aquaculture Reports. 2025;40:102626. doi:10.1016/j.aqrep.2025.102626

17. Cherbas L, Gong L. Cell lines. Methods. 2014;68(1):74–81. doi:10.1016/j.ymeth.2014.01.006

18. Lynn DE. Development and characterization of insect cell lines. Cytotechnology. 1996;20:3–11. doi:10.1007/BF00350384

19. Hesp K, Van Der Heijden JME, Munroe S, et al. First continuous marine sponge cell line established. Sci Rep. 2023;13(1):5766. doi:10.1038/s41598-023-32394-x

20. Quinn B, Costello MJ, Dorange G, Wilson JG, Mothersill C. Development of an in vitro culture method for cells and tissues from the zebra mussel (Dreissena polymorpha). Cytotechnology. 2009;59(2):121–134. doi:10.1007/s10616-009-9202-3

21. Suja CP, Sukumaran N, Dharmaraj S. Effect of culture media and tissue extracts in the mantle explant culture of abalone, Haliotis varia Linnaeus. Aquaculture. 2007;271(1-4):516–522. doi:10.1016/j.aquaculture.2007.04.086

22. Van Der Merwe M, Auzoux-Bordenave S, Niesler C, Roodt-Wilding R. Investigating the establishment of primary cell culture from different abalone (Haliotis midae) tissues. Cytotechnology. 2010;62(3):265–277. doi:10.1007/s10616-010-9293-x

23. Attaallah A, Marchionni S, El-Beltagy A, Abdelaziz K, Lorenzini A, Milani L. Cell cultures of the Manila clam and their possible use in biomonitoring and species preservation. The European Zoological Journal. 2020;87(1):624–641. doi:10.1080/24750263.2020.1827052

24. Balakrishnan S, Sajeevan AKM, Parvathi SC, Bright Singh IS, Puthumana J. An optimized protocol for routine development of cell culture from adult oyster, *Crassostrea madrasensis*. Cell Biology International. 2024;48(7):923–938. doi:10.1002/cbin.12159

25. Chen S, Wen C. Establishment of cell lines derived from oyster, Crassostrea gigas Thunberg and hard clam, Meretrix lusoria Röding. Methods in Cell Science. 1999;21:183–192. doi:10.1023/a:1009829807954

26. Potts RWA, Gutierrez AP, Cortés-Araya Y, Houston RD, Bean TP. Developments in marine invertebrate primary culture reveal novel cell morphologies in the model bivalve *Crassostrea gigas*. PeerJ. 2020;8:e9180. doi:10.7717/peerj.9180

27. Arzul I, Canier L, Chollet B, Garcia C. Bonamia spp. infections of oysters. In: Diseases of Bivalves. Elsevier; 2025:1–14. doi:10.1016/B978-0-12-820339-2.00002-4

28. Buestel D, Ropert M, Prou J, Goulletquer P. History, Status, and Future of Oyster Culture in France. Journal of Shellfish Research. 2009;28(4):813–820. doi:10.2983/035.028.0410

29. Allen H. Towards an Economic Value of Native Oyster Restoration in Scotland: Provisioning, Regulating and Cultural Ecosystem Services. Centre of Expertise for Waters; 2019.

30. Pouvreau S, Lapègue S, Arzul I, Boudry P. Fifty years of research to counter the decline of the European flat oyster (*Ostrea edulis*): a review of French achievements and prospects for the restoration of remaining beds and revival of aquaculture production. Aquat Living Resour. 2023;36:13. doi:10.1051/alr/2023006

31. Fischer C, Milting H, Fein E, et al. Long-term functional and structural preservation of precision-cut human myocardium under continuous electromechanical stimulation in vitro. Nat Commun. 2019;10(1):117. doi:10.1038/s41467-018-08003-1

32. Audino JA, Marian JEAR, Wanninger A, Lopes SGBC. Mantle margin morphogenesis in Nodipecten nodosus (Mollusca: Bivalvia): new insights into the development and the roles of bivalve pallial folds. BMC Dev Biol. 2015;15(1):22. doi:10.1186/s12861-015-0074-9

33. Mamangkey NGF, Southgate PC. Regeneration of excised mantle tissue by the silver-lip pearl oyster, Pinctada maxima (Jameson). Fish & Shellfish Immunology. 2009;27(2):164–174. doi:10.1016/j.fsi.2009.03.009

34. Chamley-Campbell J, Campbell GR, Ross R. The smooth muscle cell in culture. Physiological Reviews. 1979;59(1):1–61. doi:10.1152/physrev.1979.59.1.1

35. Eymann C, Götze S, Bock C, et al. Thermal performance of the European flat oyster, Ostrea edulis (Linnaeus, 1758)—explaining ecological findings under climate change. Mar Biol. 2020;167(2):17. doi:10.1007/s00227-019-3620-3

36. Marrec-Croq FL, Glaise D, Guguen-Guillouzo C, et al. Primary Cultures of Heart Cells from the Scallop Pecten Maximus (Mollusca-Bivalvia). In Vitro Cellular & Developmental Biology - Animal. 1999;35:289–295. doi:10.1007/s11626-999-0073-x.

37. Ballarin L, Karahan A, Salvetti A, et al. Stem Cells and Innate Immunity in Aquatic Invertebrates: Bridging Two Seemingly Disparate Disciplines for New Discoveries in Biology. Front Immunol. 2021;12:688106. doi:10.3389/fimmu.2021.688106

38. Clevers H. Modeling Development and Disease with Organoids. Cell. 2016;165(7):1586–1597. doi:10.1016/j.cell.2016.05.082

39. Haszprunar G. The molluscan rhogocyte (pore-cell, blasenzelle, cellule nucale), and its significance for ideas on nephridial evolution. J Mollus Stud. 1996;62(2):185–211. doi:10.1093/mollus/62.2.185

40. Matozzo V, Brunelli N, Cima F. The underrated immune role of bivalve ‘serous cells’: New insight from inflammatory responses of the Manila clam Ruditapes philippinarum. Fish & Shellfish Immunology. 2025;159:110188. doi:10.1016/j.fsi.2025.110188

41. Meyhofer E, Morse MP. Characterization of the Bivalve Ultrafiltration System in Mytilus edulis, Chlamys hastata, and Mercenaria mercenaria. Invertebrate Biology. 1996;115(1):20. doi:10.2307/3226939

42. Bower SM. Synopsis of Infectious Diseases and Parasites of Commercially Exploited Shellfish: Bonamia ostreae of Oysters. Published online 2015. https://www.dfo-mpo.gc.ca/science/aah-saa/diseases-maladies/bonostoy-eng.html

43. Bigas M, Durfort M, Poquet M. Cytological effects of experimental exposure to Hg on the gill epithelium of the European flat oyster Ostrea edulis: ultrastructural and quantitative changes related to bioaccumulation. Tissue and Cell. 2001;33(2):178–188. doi:10.1054/tice.2000.0169

44. Roopnarine DS, Roopnarine PD, Anderson LC, Hwang JH, Patel S. Metaplasia of respiratory and digestive tissues in the eastern oyster Crassostrea virginica associated with the Deepwater Horizon oil spill. Sanganyado E, ed. PLoS ONE. 2021;16(9):e0247739. doi:10.1371/journal.pone.0247739

45. Rock JR, Hogan B. Epithelial Progenitor Cells in Lung Development, Maintenance, Repair, and Disease. Annual review of cell and developmental biology. 2011;27. doi:10.1146/annurev-cellbio-100109-104040

46. Fernández A, Câmara N, Sierra E, et al. Cetacean Intracytoplasmic Eosinophilic Globules: A Cytomorphological, Histological, Histochemical, Immunohistochemical, and Proteomic Characterization. Animals. 2023;13(13):2130. doi:10.3390/ani13132130

47. Zatloukal K, French SW, Stumptner C, et al. From Mallory to Mallory–Denk bodies: What, how and why? Experimental Cell Research. 2007;313(10):2033–2049. doi:10.1016/j.yexcr.2007.04.024

48. Machii A. Organ culture of mantle tissue of the pearl oyster Pinctada fucata (Gould). Bulletin of the National Pearl Research Laboratory. 1974;18:2111–2117.

49. Awaji M, Machii A. Fundamental studies on in vivo and in vitro pearl formation—Contribution of outer epithelial cells of pearl oyster mantle and pearl sacs. Aqua-BioScience Monographs. 2011;4(1):1–39. doi:10.5047/absm.2011.00401.0001

50. De La Forest Divonne S, Pouzadoux J, Romatif O, et al. Diversity and functional specialization of oyster immune cells uncovered by integrative single-cell level investigations. eLife. 2025;13:RP102622. doi:10.7554/eLife.102622

51. Rebelo MDF, Figueiredo EDS, Mariante RM, Nóbrega A, De Barros CM, Allodi S. New Insights from the Oyster Crassostrea rhizophorae on Bivalve Circulating Hemocytes. Singh SR, ed. PLoS ONE. 2013;8(2):e57384. doi:10.1371/journal.pone.0057384

52. Checa AG. Physical and Biological Determinants of the Fabrication of Molluscan Shell Microstructures. Front Mar Sci. 2018;5:353. doi:10.3389/fmars.2018.00353

53. Xiang L, Kong W, Su J, et al. Amorphous Calcium Carbonate Precipitation by Cellular Biomineralization in Mantle Cell Cultures of Pinctada fucata. Strack S, ed. PLoS ONE. 2014;9(11):e113150. doi:10.1371/journal.pone.0113150

54. Ivanina AV, Falfushynska HI, Beniash E, Piontkivska H, Sokolova IM. Biomineralization-related specialization of hemocytes and mantle tissues of the Pacific oysters *Crassostrea gigas*. Journal of Experimental Biology. Published online January 1, 2017:jeb.160861. doi:10.1242/jeb.160861

55. Sillanpää JK, Ramesh K, Melzner F, Sundh H, Sundell K. Calcium mobilisation following shell damage in the Pacific oyster, Crassostrea gigas. Marine Genomics. 2016;27:75–83. doi:10.1016/j.margen.2016.03.001

56. Lian S, Hu N, Chen X, et al. Widespread presence of bone marrow–like hematopoietic stem cell niche in invertebrate skeletons. Science AdvAnceS. Published online 2025.

57. Song X, Liu Z, Wang L, Song L. Recent Advances of Shell Matrix Proteins and Cellular Orchestration in Marine Molluscan Shell Biomineralization. Front Mar Sci. 2019;6:41. doi:10.3389/fmars.2019.00041

58. Dessai SN. Primary culture of mantle cells of bivalve mollusc, Paphia malabarica. In Vitro CellDevBiol-Animal. 2012;48(8):473–477. doi:10.1007/s11626-012-9538-4

59. Endoh M, Hasegawa Y. Culture of mantle epithelial cells expressing shell matrix proteins from scallop Patinopecten yessoensis. Fisheries Sci. 2006;72(6):1277–1285. doi:10.1111/j.1444-2906.2006.01286.x

